# The flavin-containing monooxygenases OSGIN1 and OSGIN2 antagonize each other to favor cytokinesis completion

**DOI:** 10.64898/2026.09.08.750123

**Authors:** Léa Lacroix, Eugénie Goupil, Matthew J. Smith, Jean-Claude Labbé

## Abstract

Cytokinesis is a fundamental process that bisects a mother cell into two distinct daughter cells via an ingressing ring of actomyosin. It initiates after anaphase chromosome separation by the mitotic spindle, which coordinates the activation of the small GTPase RhoA at the cell equator. We previously identified OSGIN1 as a flavin-containing monooxygenase enzyme required for cytokinesis completion via the regulation of RhoA activity. Here we show that the OSGIN1 paralog OSGIN2 is also a flavin-containing monooxygenase implicated in cytokinetic regulation. Cells lacking OSGIN2 have reduced RhoA activity and increased rates of cytokinesis failure, similar to *OSGIN1*-deleted cells. Strikingly, these phenotypes are suppressed in cells lacking both OSGIN1 and OSGIN2, indicating that the two proteins antagonize each other. Consistent with this, we find that OSGIN1 and OSGIN2 form both homo- and heteromeric complexes *in vitro* and in cells. Together, our results support a model in which OSGIN1 and OSGIN2 both function as negative regulators of RhoA signaling and that their heteromeric interaction during mitosis antagonizes their activity to favor cytokinesis completion.

## Introduction

Cytokinesis is the last stage of cell division and separates a mother cell into two daughters. It initiates after anaphase onset via the assembly of an actomyosin ring at the cell equator that pulls the plasma membrane towards the spindle center as it ingresses, effectively separating the duplicated sets of chromosomes into distinct entities (reviewed in Green et al., 2012). In early cytokinesis, assembly and regulation of the actomyosin ring depend on the cortical activation of the small GTPase RhoA which, upon GTP binding, locally engages downstream effectors that favor actin polymerization and myosin activity (Basant and Glotzer, 2018; Green et al., 2012). RhoA activity is required throughout cytokinesis, including in late cytokinesis when it enables the transient stabilization of the intercellular bridge around the midbody, the organelle that coordinates the final membrane scission (Andrade and Echard, 2022; Green et al., 2012).

Previous work has revealed that *C. elegans* OSGN-1 and human OSGIN1 are cytokinetic regulators (Goupil et al., 2024). These proteins are flavin-containing monooxygenases (FMOs), a class of enzymes that employ flavin adenine dinucleotide (FAD) and reduced nicotinamide adenine dinucleotide phosphate (NADPH) as cofactors to catalyze the transfer of an oxygen atom onto a substrate (Huijbers et al., 2014; Lacroix et al., 2024; Rossner et al., 2017). In HeLa cells, OSGIN1 was found to localize at the intercellular bridge in late cytokinesis and its deletion decreased the levels of active RhoA at the bridge, leading to defects in bridge stability and a significant increase in the proportion of multinucleated cells. These phenotypes could be rescued by exogenously expressing either *C. elegans* OSGN-1 or human OSGIN1, establishing these proteins as functional orthologs required for intercellular bridge stability in late cytokinesis (Goupil et al., 2024).

The human OSGIN1 paralog, OSGIN2, shares a similar architecture, with conserved domains bearing high amino acid identity at key positions (Figure 1A-B). Deletion of *OSGIN2* in HeLa cells was also found to increase the proportion of multinucleated cells. However, unlike *OSGIN1* deletion, the defect could not be rescued by expressing *C. elegans* OSGN-1 (Goupil et al., 2024). This suggests that the function of OSGIN2 is not redundant with that of OSGIN1, leaving the mechanism by which it impacts HeLa cell ploidy unresolved.

**Figure 1.**
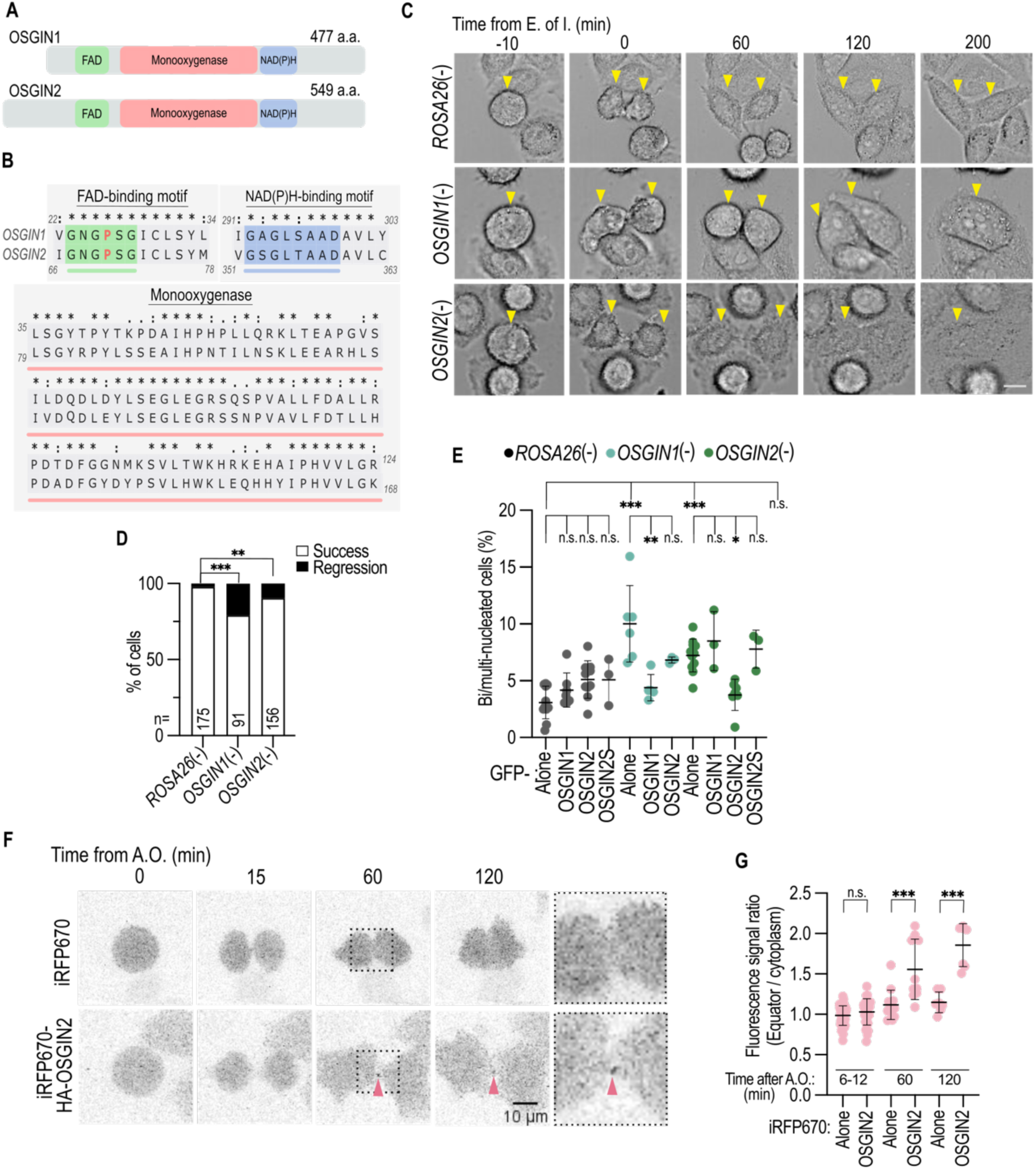
OSGIN2 regulates cytokinesis. (A) Schematic organization of OSGIN1 and OSGIN2 depicting the presumptive position of the FAD-binding, monooxygenase and NADPH-binding domains. (B) Amino acid sequence alignment of OSGIN1 and OSGIN2 domains showing the two Rossman folds (FAD- and NADPH-binding) and the monooxygenase region. A conserved proline (in red) required for activity is highlighted in the FAD-binding motif of both protein sequences. (C-D) Time-lapse brightfield images of dividing *ROSA26*(-) control, *OSGIN1*(-) and *OSGIN2*(-) HeLa cells (C) and quantification of cytokinetic outcome in the same conditions (D). Time (in min) is relative to the end of furrow ingression (E. of I.). Arrowheads denote the dividing mother cell and daughter cells. Scale bar = 10 µm. n = 91-215 cells acquired in N = 3 replicates. n.s. = not significant, **p<0.01, \*\*\**p* < 0.001, Fisher’s exact test. (E) Quantification of the proportion of bi/multinucleated *ROSA26*(-) control, *OSGIN1*(-) and *OSGIN2*(-) HeLa cells stably expressing either GFP (alone), GFP-OSGIN1, GFP-OSGIN2 or GFP-OSGIN2S. n=72-919 cells scored in N ≥ 3 replicates. Bars denote mean ± SD. n.s. = not significant, **p<0.01, *p<0.05, \*\**p* < 0.01, \*\*\**p* < 0.001, one-way ANOVA with Tukey’s multiple comparison correction. See Figure 2A for representative images. (F-G) Time-lapse confocal images (F) and quantification of the average ratio of equator/cytoplasm iRFP670 fluorescence signal (G) of control cells expressing either iRFP670 (alone) or iRFP670-OSGIN2. Arrowheads in (F) point to the enriched signal at the intercellular bridge. Time (in min) is relative to anaphase onset (A.O.). n = 7-53 cells scored in N ≥ 3 replicates. Bars denote mean ± SD. n.s. = not significant, \*\*\**p* < 0.001, one-way ANOVA with Sidak’s multiple comparison correction. Scale bar = 10 µm, with the boxed region at 60 min magnified 3X in insets.

Here, we show that OSGIN2 is also an FMO that regulates RhoA activity during cytokinesis and is required for cytokinetic completion. Interestingly, we find that while inactivating OSGIN1 or OSGIN2 individually results in intercellular bridge stability defects and cytokinetic furrow regression, the simultaneous deletion of both genes suppresses these defects and results in an increase in RhoA activity, denoting an antagonistic relationship between the two human paralogs. Accordingly, we find that OSGIN1 and OSGIN2 form both homo- and heteromeric complexes. Our work suggests a model in which OSGIN1 and OSGIN2 redundantly limit RhoA activity during cytokinesis and antagonize each other in late cytokinesis to stabilize the intercellular bridge and ensure cytokinetic furrow stability.

## Results

### OSGIN2 is a late cytokinesis regulator

To determine whether OSGIN2 impacts HeLa cell ploidy by regulating cytokinesis, we performed live imaging of *OSGIN2*-deleted HeLa cells undergoing mitosis and scored the proportion of cells that failed to complete final membrane scission (termed abscission). We found that all cells initiated furrow ingression normally, indicating that OSGIN2 activity is not required in early cytokinesis. However a significant proportion of *OSGIN2*-deleted cells underwent furrow regression in late cytokinesis, similar to what was observed in *OSGIN1*-deleted cells (Figure 1C-D). Furthermore, we found that deleting *OSGIN2* in HEK 293 cells resulted in an increase in the proportion of multinucleated cells (Figure S1), indicating that the requirement for OSGIN2 is not specific to HeLa cells and thus that OSGIN2 is generally required for cytokinesis.

The *OSGIN2* locus is predicted to encode two isoforms, a canonical one of 61 kDa (OSGIN2) and a 56 kDa isoform with shorter N-terminal region (hereafter OSGIN2S), both of which were compromised by the CRISPR/Cas9-mediated deletion strategy. To assess if one or both isoforms are required for cytokinesis, we re-expressed them individually in *OSGIN2*-deleted HeLa cells and assessed if they could rescue cytokinesis defects by scoring the proportion of multinucleated cells. We found that re-expressing OSGIN2, but not OSGIN2S, completely rescued HeLa cell multinucleation (Figure 1E), indicating that only the canonical isoform of OSGIN2 functions in cytokinesis. Consistent with a role in late cytokinesis, live-imaging of HeLa cells expressing an iRFP-tagged version of OSGIN2 revealed that it localizes at the intercellular bridge (Figure 1F-G), where OSGIN1 was also found to be enriched (Goupil et al., 2024). Together, these results indicate that OSGIN2, like OSGIN1, is a late cytokinesis regulator.

### OSGIN2 is a flavin-containing monooxygenase and regulates RhoA activity

Both human OSGIN1 and its *C. elegans* ortholog, OSGN-1, bind to FAD and have monooxygenase activity *in vitro*, and regulate the activity of the small GTPase RhoA *in vivo* (Goupil et al., 2024). Furthermore, replacing a conserved proline in the FAD-binding domain of *C. elegans* OSGN-1 with a leucine abrogates FAD binding and catalytic activity, and compromises the sustained activity of RhoA at the intercellular bridge, resulting in bridge stability defects (Goupil et al., 2024). We sought to determine whether OSGIN2 has monooxygenase activity *in vitro,* but the protein was largely insoluble when produced in *E. coli*. To determine whether OSGIN2 possesses catalytic activity, we therefore introduced an analogous proline-to-leucine inactivating mutation in its FAD-binding domain (Figure 1B) and assessed the ability of this OSGIN2^P70L^ variant to rescue the HeLa cell ploidy defect resulting from the loss of *OSGIN2*. We found that the multinucleation defect of *OSGIN2*-deleted cells was rescued by expressing wild-type OSGIN2 but not the OSGIN2^P70L^ variant (Figure 2A-B). Similar results were obtained in *OSGIN1*-deleted HeLa cells, in which the multinucleation defect was rescued by expressing wild-type OSGIN1 but not the OSGIN1^P26L^ variant (Figure 2A-B). Immunofluorescence analysis revealed that both OSGIN1^P26L^ and OSGIN2^P70L^ variants localized at the intercellular bridge in late HeLa cell cytokinesis (Figure 2C), consistent with the two proteins being expressed and trafficked as in control cells. These results indicate that OSGIN2, like human OSGIN1 and *C. elegans* OSGN-1, is an FMO and that its activity is required in late cytokinesis. Tracking the localization and dynamics of an *in vivo* RhoA activity sensor (the Rho-binding domain of Rhotekin fused to dTomato; Mahlandt et al., 2021) previously revealed that OSGIN1 is required to maintain RhoA activity during cytokinesis (Goupil et al., 2024) (Figure 3A-C). To determine whether OSGIN2 also modulates RhoA activity during cytokinesis, we measured the levels of this RhoA sensor in *OSGIN2*-deleted HeLa cells. We found that deletion of *OSGIN2* resulted in a small, albeit non-statistically significant, decrease of RhoA sensor levels at the cytokinetic furrow and intercellular bridge compared to the control (Figure 3A-C). Distribution of the RhoA sensor nevertheless appeared compromised in *OSGIN2*-deleted HeLa cells, which displayed an increased variation of sensor levels compared to both control and *OSGIN1*-deleted cells (Figure 3C). Furthermore, the observed decrease in RhoA sensor levels correlates with the lower impact on cytokinesis completion of deleting *OSGIN2* compared to *OSGIN1* (Figure 1D). These results indicate that while OSGIN1 is required to sustain RhoA activity in late cytokinesis, the impact of OSGIN2 on RhoA activity may not be as significant.

**Figure 2.**
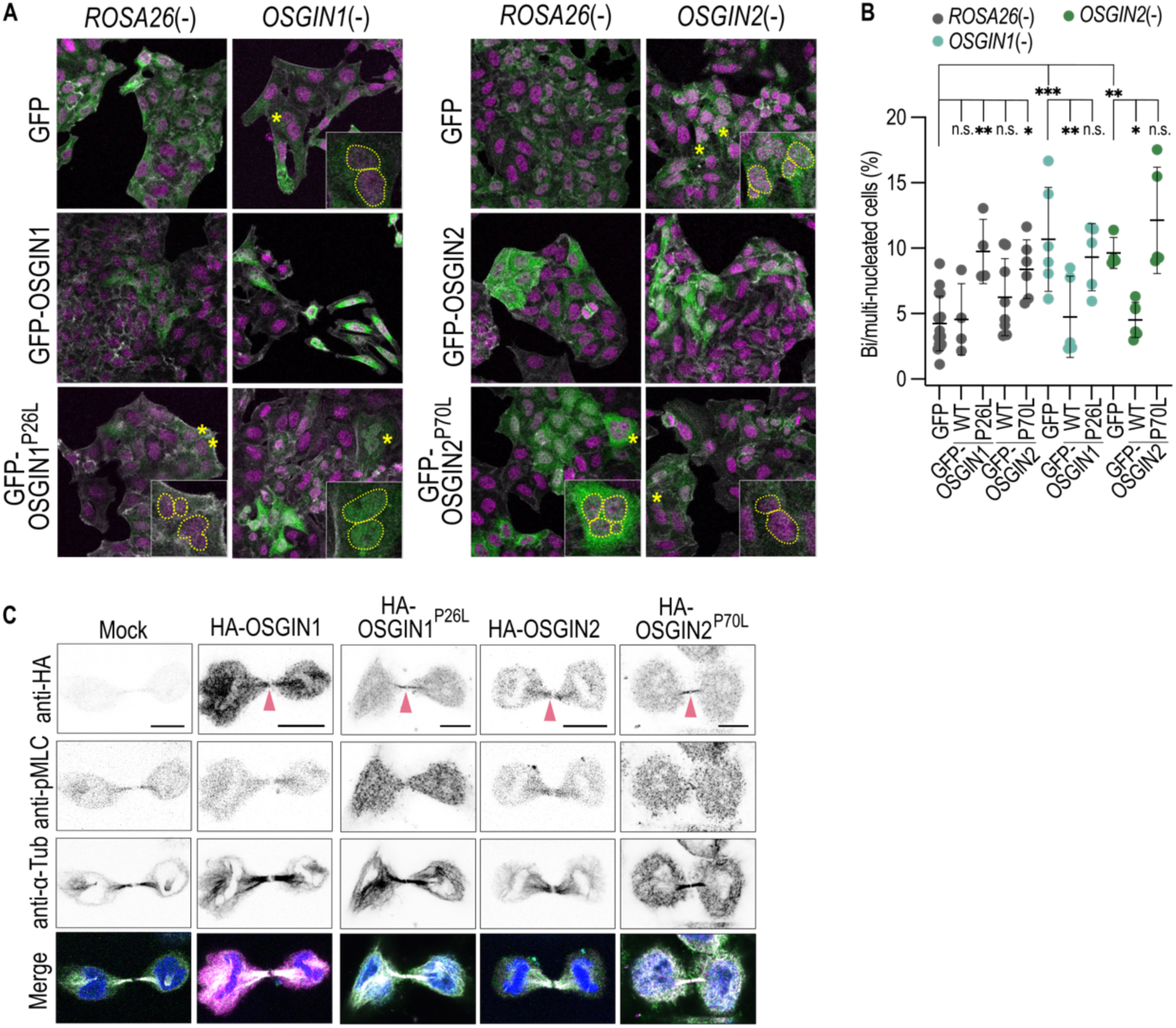
The flavin-containing monooxygenase activity of OSGIN2 is required for cytokinesis. (A) Representative indirect immunofluorescence images of *ROSA26*(-) control or *OSGIN1*(-) (left panels) and *ROSA26*(-) control or *OSGIN2*(-) (right panel) HeLa cells stably expressing either, GFP, GFP-OSGIN1 (WT or P^26^L) or GFP-OSGIN2 (WT or P^70^L), as indicated. Cells were stained with anti-GFP (green), phalloidin (revealing F-actin, white) and Hoechst 33342 (revealing nuclei, magenta). Yellow asterisks denote bi/multinucleated cells, magnified 4 times in insets, in which nuclei are outlined. Scale bar = 50 µm. (B) Quantification of the proportion of bi/multinucleated *ROSA26*(-) control, *OSGIN1*(-) and *OSGIN2*(-) HeLa cells stably expressing either GFP, GFP-OSGIN1 (WT or P^26^L), or GFP-OSGIN2 (WT or P^70^L), as indicated. n=78-295 cells scored in N ≥ 3 replicates. Bars denote mean ± SD. n.s. = not significant, \**p* < 0.05, \*\**p* < 0.01, \*\*\**p* < 0.001, one-way ANOVA with Sidak’s multiple comparison correction. (C) Representative indirect immunofluorescence images of control HeLa cells in telophase either untransfected (mock) or transiently expressing HA-OSGIN1 (WT or P^26^L) or HA-OSGIN2 (WT or P^70^L). Cells were stained with anti-HA (magenta), anti-phospho-MLC (green), anti-α-tubulin (white) and Hoechst 33342 (revealing nuclei, blue). Arrowheads in the HA stains (top row) point to the intercellular bridge. Scale bars = 10 µm.

**Figure 3.**
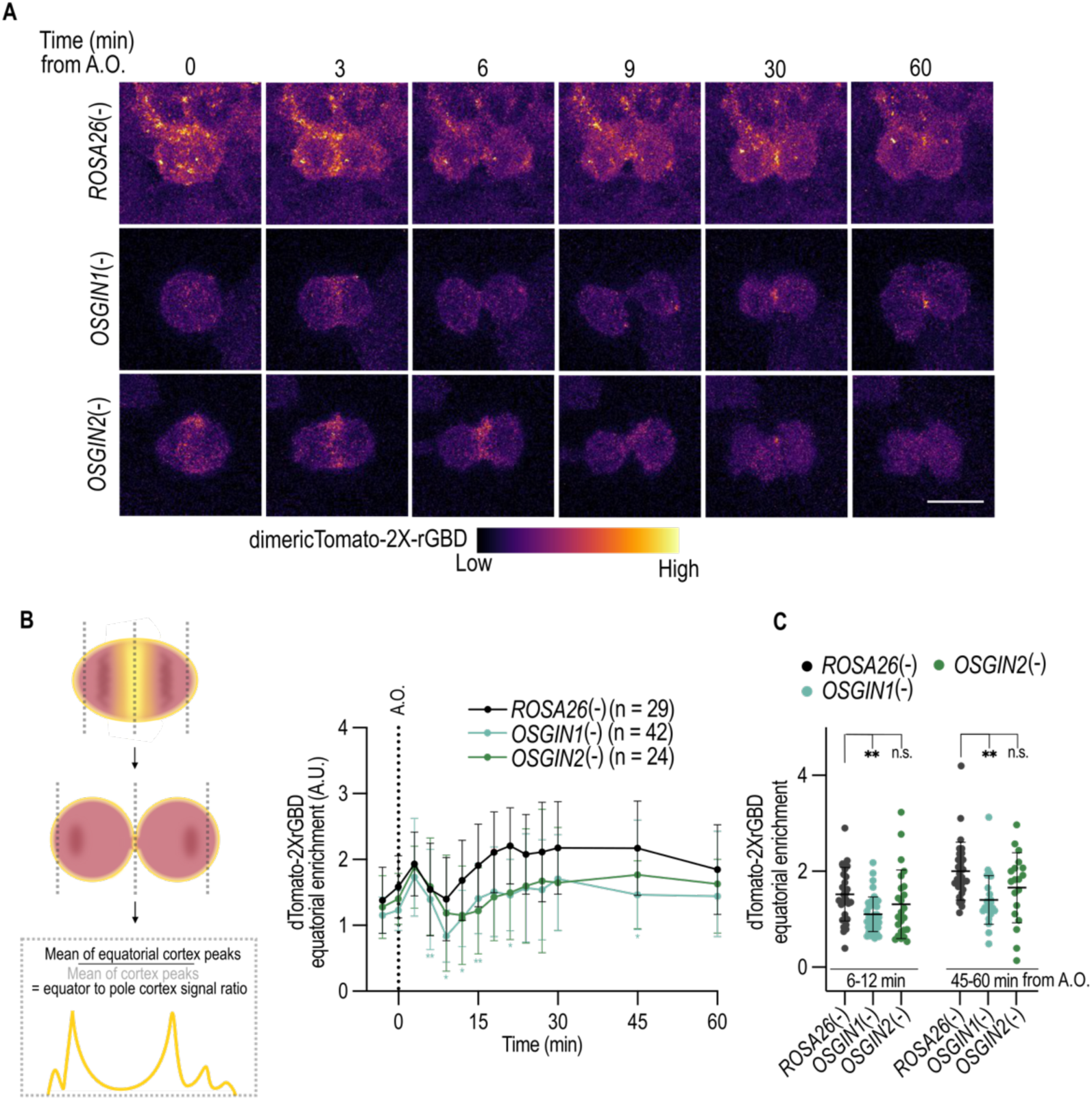
OSGIN1 and OSGIN2 regulate RhoA activity in late cytokinesis. (A-B) Time-lapse confocal images (A) and measurement of equator/cortex fluorescence levels over time (B) of the dimericTomato-2X-rGBD RhoA activity reporter (colored scale) stably expressed in *ROSA26*(-) control, *OSGIN1*(-) or *OSGIN2*(-) HeLa cells. The schematic illustration in (B) depicts the fluorescence quantification method. Values correspond to the mean ± SD over three or more biological replicates (n = 24-42 cells per condition, as indicated). n.s. = not significant, \*\**p* < 0.01, \*\*\**p* < 0.001, two-way ANOVA with Sidak’s multiple comparison correction. Time (in min) is relative to anaphase onset (A.O.). Scale bar = 20 µm. (C) Average fluorescence intensity levels of the dimericTomato-2X-rGBD reporter pooled in early (6-12 min after A.O.) and late (45-60 min after A.O.) cytokinesis. Bars indicate mean ± SD. ns = not significant, \**p* < 0.05, \*\**p* < 0.01, one-way ANOVA with Tukey’s multiple comparison correction.

### OSGIN1 and OSGIN2 antagonize each other’s activity in late cytokinesis

These results, together with our previous finding that expressing human OSGIN1 or *C. elegans* OSGN-1 in *OSGIN2*-deleted HeLa cells did not rescue their multinucleation defects (Goupil et al., 2024), suggest that OSGIN1 and OSGIN2 are not redundant. To test this, we first performed cross-rescue experiments by overexpressing one gene in cells mutant for the other gene and quantified the impact on ploidy. We found that while expression of OSGIN1 in *OSGIN1*-deleted cells rescued the multinucleation defects, expressing OSGIN2 in *OSGIN1*-deleted cells had no impact on their ploidy (Figure 1E). Likewise, expressing OSGIN2, but not OSGIN1, in *OSGIN2*-deleted cells decreased the proportion of multinucleated cells (Figure 1E). These results indicate that OSGIN1 and OSGIN2 cannot compensate each other’s function in HeLa cell cytokinesis.

To further explore the individual role of the two proteins, we employed CRISPR/Cas9 to simultaneously delete both *OSGIN1* and *OSGIN2* and assessed the impact on HeLa cell ploidy. Strikingly, we found that deletion of both *OSGIN1* and *OSGIN2* (hereafter *OSGIN1/2*-deleted) restored normal rates of cytokinesis completion (Figure 4A-B) and resulted in a proportion of multinucleated cells that is similar to control, significantly decreased compared to cells individually deleted for either gene (Figure 4C-D). A similar impact on ploidy was found in HEK 293 cells, in which depleting OSGIN1 by siRNA treatment in *OSGIN2*-deleted cells restored normal ploidy compared to either mock-depleted *OSGIN2*-deleted cells or *OSGIN1*-depleted control cells (Figure S1). This phenotypic suppression was specific to the loss of the two genes, as expressing either OSGIN1 or OSGIN2 in *OSGIN1/2*-deleted HeLa cells resulted in an increase in multinucleation defects (Figure 4C-D), partially recapitulating what is observed in either single mutant. This increase was not observed when expressing the catalytically-inactive OSGIN1^P26L^ or OSGIN2^P70L^ variants (Figure 4D), indicating that the monooxygenase activity of the enzymes is needed in this context. These results indicate that OSGIN1 and OSGIN2 antagonize each other’s activity in late cytokinesis and further suggest that the ploidy defects observed in each single mutant arise because of the misregulated activity of the other OSGIN protein.

**Figure 4.**
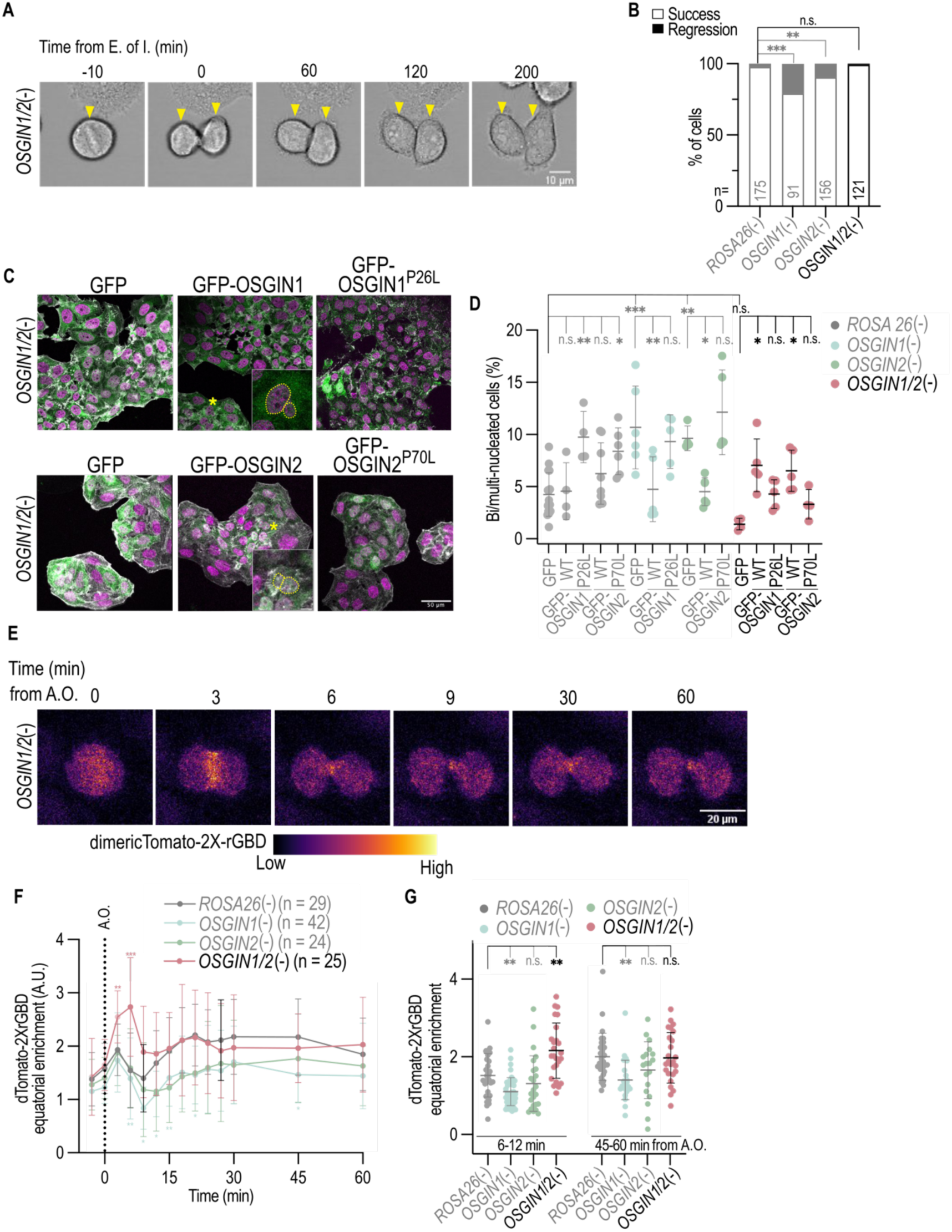
OSGIN1 and OSGIN2 antagonize each other’s activity during cytokinesis. (A-B) Time-lapse brightfield images of dividing *OSGIN1/2*(-) HeLa cells (A) and quantification of cytokinetic outcome in *ROSA26*(-) control, *OSGIN1*(-), *OSGIN2*(-) and *OSGIN1/2*(-) cells (B). Time (in min) is relative to the end of furrow ingression (E. of I.). Arrowheads denote the dividing mother cell and daughter cells. Scale bar = 10 µm. n = 91-215 cells acquired in N = 3 replicates. n.s. = not significant, **p<0.01, \*\*\**p* < 0.001, Fisher’s exact test. (C) Representative indirect immunofluorescence images of *OSGIN1/2*(-) HeLa cells stably expressing either, GFP, GFP-OSGIN1 (WT or P^26^L) or GFP-OSGIN2 (WT or P^70^L), as indicated. Cells were stained with anti-GFP (green), phalloidin (revealing F-actin, white) and Hoechst 33342 (revealing nuclei, magenta). Yellow asterisks denote bi/multinucleated cells, magnified 4 times in insets, in which nuclei are outlined. Scale bar = 50 µm. (D) Quantification of the proportion of bi/multinucleated *ROSA26*(-) control, *OSGIN1*(-), *OSGIN2*(-) and *OSGIN1/2*(-) HeLa cells stably expressing either GFP, GFP-OSGIN1 (WT or P^26^L), or GFP-OSGIN2 (WT or P^70^L), as indicated. n=78-295 cells scored in N ≥ 3 replicates. Bars denote mean ± SD. n.s. = not significant, \**p* < 0.05, \*\**p* < 0.01, \*\*\**p* < 0.001, one-way ANOVA with Sidak’s multiple comparison correction. (E-F) Time-lapse confocal images of *OSGIN1/2*(-) HeLa cells (E) and measurement of equator/cortex fluorescence levels over time (F) of the dimericTomato-2X-rGBD RhoA activity reporter (colored scale) stably expressed in *ROSA26*(-) control, *OSGIN1*(-), *OSGIN2*(-) or *OSGIN1/2*(-) HeLa cells. Values correspond to the mean ± SD over three or more biological replicates (n = 24 to 42 cells in total per condition, as indicated). n.s. = not significant, \*\**p* < 0.01, \*\*\**p* < 0.001, two-way ANOVA with Sidak’s multiple comparison correction. Time (in min) is relative to anaphase onset (A.O.). Scale bar = 20 µm. (G) Average fluorescence intensity levels of the dimericTomato-2X-rGBD reporter pooled in early (6-12 min after A.O.) and late (45-60 min after A.O.) cytokinesis. Bars indicate mean ± SD. ns = not significant, \**p* < 0.05, \*\**p* < 0.01, one-way ANOVA with Tukey’s multiple comparison correction. In panels B, E, G and H, the represented data for *ROSA26*(-) control, *OSGIN1*(-) and *OSGIN2*(-) conditions is respectively reproduced from Figures 1D, 2B, 3B and 3C, to help comparisons, and faded to highlight the data for *OSGIN1/2(-)* conditions.

### OSGIN1 and OSGIN2 redundantly limit RhoA activity in early cytokinesis

Our finding that OSGIN1 and OSGIN2 antagonize each other to favor cytokinesis completion suggested that each OSGIN protein redundantly regulates some aspects of cytokinesis. To assess this, we employed live imaging to track the dynamics of HeLa cell division in control and *OSGIN* mutant cells. We first compared the dynamics of cytokinetic furrow ingression between the various conditions by monitoring mRFP fused to a CAAX motif, enabling the tracking of membrane closure at the cell equator over time (Figure 5A). We found that cytokinetic furrow ingression occurs with a comparable rate in all conditions (Figure 5B-C), suggesting that the composition and regulation of the cytokinetic ring is similar. However, *OSGIN1/2*-deleted cells initiated furrow ingression significantly earlier after anaphase onset than control cells or cells individually deleted for either *OSGIN1* or *OSGIN2* (Figure 5B). This suggests that OSGIN1 and OSGIN2 redundantly limit the timing and/or rate of actomyosin ring assembly without impacting the rates cytokinetic furrow ingression *per se*.

**Figure 5.**
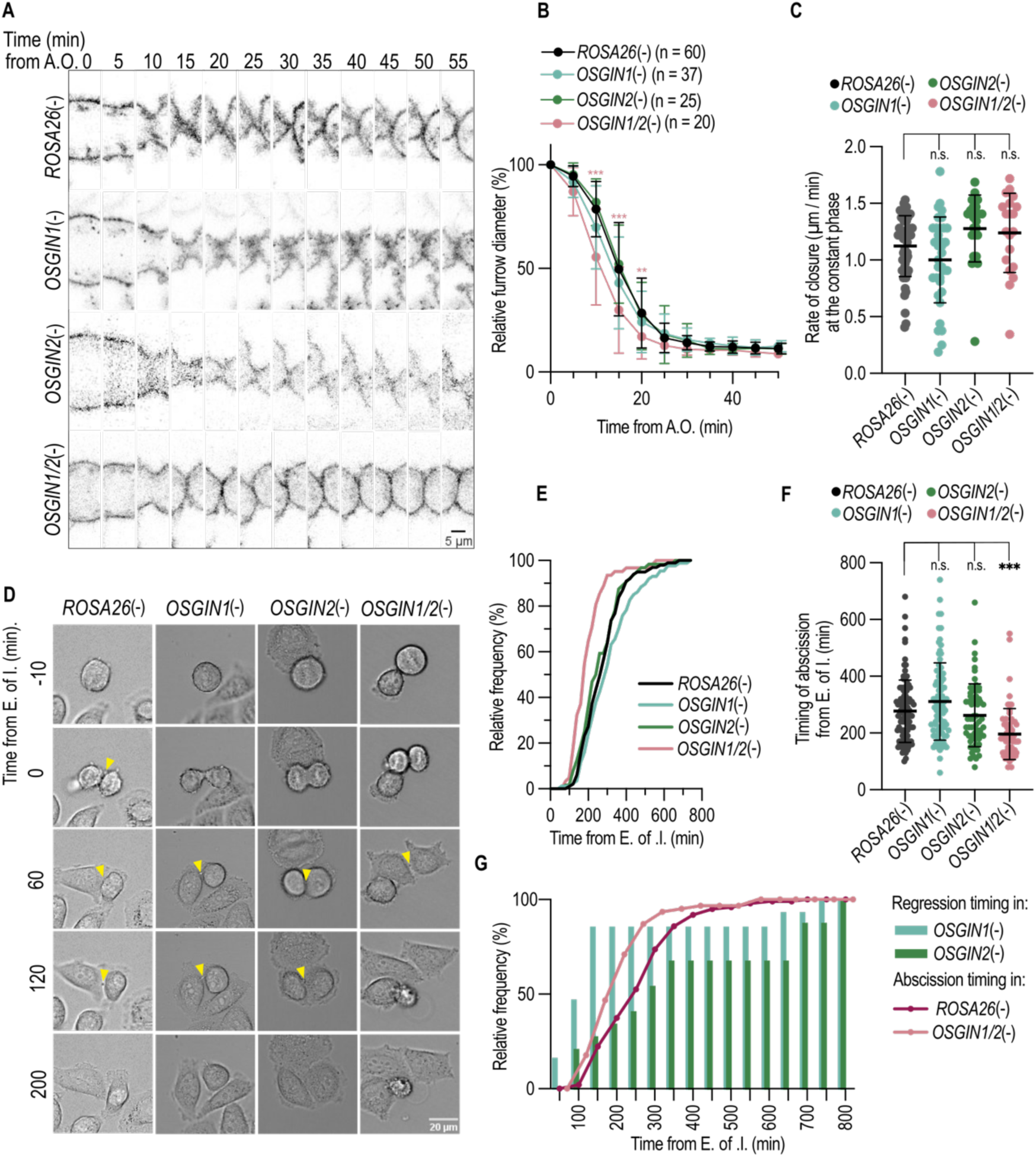
OSGIN1 and OSGIN2 impact cytokinesis progression and completion. (A) Time-lapse confocal images (single slice) at the equatorial region of dividing *ROSA26*(-) control, *OSGIN1*(-), *OSGIN2*(-) and *OSGIN1/2*(-) cells expressing the mRFP-CAAX membrane reporter. Time (in min) is relative to anaphase onset (A.O.). Scale bar = 5 µm. (B-C) Quantification of the change in cytokinetic furrow diameter over time (in min) relative to that at anaphase onset (B) and quantification of the rate of cytokinetic furrow ingression (C) at the constant phase (in µm/min) in the same conditions as in (A). Bars indicate mean ± SD. n=20-60 cells acquired in N>3 replicates. n.s. = not significant, \*\**p* < 0.01, \*\*\**p* < 0.001, two-way ANOVA with repeated measures and Dunnett’s multiple comparison correction (B) or one-way Anova with Dunnett’s multiple comparison correction (C). (D) Time-lapse brightfield images of dividing *ROSA26*(-) control, *OSGIN1*(-), *OSGIN2*(-) and *OSGIN1/2*(-) HeLa cells. Time (in min) is relative to the end of furrow ingression (E. of I.). Arrowheads denote the intercellular bridge prior to abscission. Scale bar = 20 µm. (E-F) Cumulative frequency of cells completing abscission over time (E) and quantification of the timing of abscission (in min) after E. of I. (F). n=57-72 cells acquired in N=3 replicates. Bars in (F) denote mean ± SD. n.s. = not significant, \*\*\**p* < 0.001, one-way ANOVA with Dunnett’s multiple comparison correction. (G) Cumulative frequencies of cytokinetic furrow regression (cytokinetic failure) over time in *OSGIN1*(-) (n=13) and *OSGIN2*(-) (n=15) HeLa cells, and of cells completing abscission over time in *ROSA26*(-) control (n=99) and *OSGIN1/2*(-) (n=62) HeLa cells, all relative to the E. of I. The furrow in *OSGIN1*(-) cells regresses prior to abscission in *OSGIN1/2*(-) cells.

We next tracked the dynamics of events at the end of mitosis, as cells complete abscission. As a certain proportion of cells deleted for *OSGIN1* or *OSGIN2* undergo cytokinetic failure, we only scored cells that completed cytokinesis during the acquisition. Live brightfield imaging revealed that *OSGIN1/2*-deleted cells underwent final membrane scission significantly earlier than control cells and cells individually deleted for either *OSGIN1* or *OSGIN2* (Figure 5D-F). Tracking the fluorescence levels of Citrin-labelled α-1B-tubulin at the intercellular bridge revealed that the timing and dynamics of microtubule clearance are comparable in all four conditions (Figure S2), indicating that the impact of OSGIN1 and OSGIN2 on the timing of abscission is independent of microtubule midbody processing. These results suggest that OSGIN1 and OSGIN2 redundantly regulate the timing of abscission at the end of mitosis. Interestingly, comparing the timing of abscission in *OSGIN1/2*-deleted cells with the timing of cytokinetic furrow regression in single mutants revealed that *OSGIN1* mutants start to undergo furrow regression before *OSGIN1/2*-deleted cells and control cells initiate abscission (Figure 5G). This suggests that furrow regression in *OSGIN1* mutant cells is a consequence of the misregulated activity of OSGIN2.

To further assess the antagonistic relationship between OSGIN1 and OSGIN2, we employed live imaging to track the dynamics of the RhoA activity sensor in *OSGIN1/2*-deleted HeLa cells. As expected based on the suppression of cytokinesis failure and ploidy defects that we observed, we found that the levels of the sensor at the intercellular bridge of *OSGIN1/2*-deleted cells were comparable to those of control cells and higher than those of either single mutant (Figure 4E-G). Strikingly however, while deleting *OSGIN2* alone had no effect on the equatorial levels of the RhoA activity sensor at the onset of cytokinetic furrow ingression (6-12 minutes after anaphase onset) and deleting *OSGIN1* alone caused a decrease in sensor levels, we found that the levels of the sensor were significantly increased at this time in *OSGIN1/2*-deleted cells (Figure 4F-G). Monitoring sensor distribution during this time also revealed that it is found in a narrower cortical band in *OSGIN1/2*-deleted cells compared to control cells and cells individually deleted for either *OSGIN1* or *OSGIN2* (Figure S3). Altogether, these results support the notion that OSGIN1 and OSGIN2 antagonize each other throughout cytokinesis and that they redundantly limit RhoA activity at the onset of furrow ingression.

### OSGIN1 and OSGIN2 can form homo- and heteromers

The antagonistic functional relationship between OSGIN1 and OSGIN2 during cytokinesis suggested that they could physically interact with one another. In support of this view, we found that overexpression of either OSGIN1^P26L^ or OSGIN2^P70L^ variant results in an increase of multinucleation in control cells (which endogenously express both proteins) but not in *OSGIN1/2*-deleted cells (Figure 4C-D). To investigate the possibility of a physical interaction, we sought to determine if these proteins can form a complex. We co-expressed GFP-HA- and TagBFP-3xFLAG-tagged versions of OSGIN1 and OSGIN2 and conducted immunoprecipitations from HEK 293 cell extracts using anti-FLAG antibodies. We found that both GFP-HA-OSGIN1 and GFP-HA-OSGIN2 were co-precipitated from cell extracts with either TagBFP-3xFLAG-OSGIN1 or TagBFP-3xFLAG-OSGIN2 (Figure 6A), consistent with OSGIN1 and OSGIN2 interacting in both homo- and heteromeric complexes. These interactions are unlikely to require catalytic activity, as GFP-HA-OSGIN1^P26L^ and GFP-HA-OSGIN2^P70L^ were likewise co-immunoprecipitated under the same conditions (Figure 6A). Furthermore, we found that *E. coli*-purified, 6xHis-tagged OSGIN1 could associate with a purified, MBP-tagged OSGIN1 but not MBP alone (Figure 6B), consistent with a direct interaction. These results suggest that OSGIN1 and OSGIN2 can form both homo- and heteromeric complexes and further indicate that OSGIN1 can directly homodimerize.

**Figure 6.**
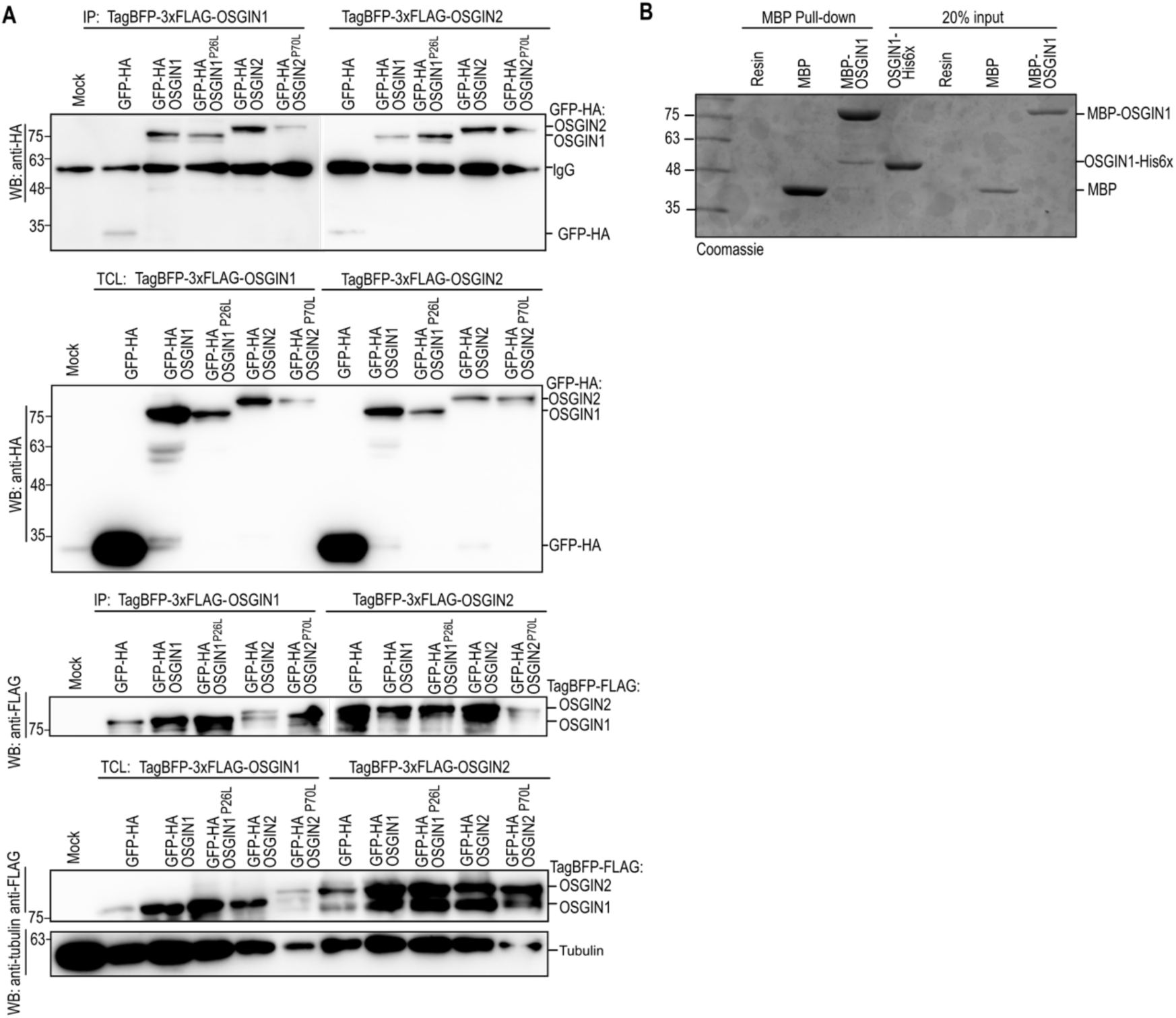
OSGIN1 and OSGIN2 form homo- and heteromeric complexes *in vitro.* (A) Western blot (WB) analyses of anti-FLAG immunoprecipitations (IP) from HeLa cells untransfected (mock) or co-expressing GFP-HA, GFP-HA-OSGIN1 (WT or P^26^L), or GFP-HA-OSGIN2 (WT or P^70^L) with either TagBFP-3xFLAG-OSGIN1 or TagBFP-3xFLAG-OSGIN2, and revealed with anti-HA (top) or anti-FLAG (bottom) antibodies, as indicated. Western blot analyses of total cell lysates (TCL) were conducted on 10% of the amounts used in the immunoprecipitations. Tubulin levels were used as loading control. (B) Coomassie staining of SDS-PAGE showing MBP (amylose resin) pull down of bacterially-purified OSGIN1-6xHis with bacterially-purified MBP-6xHis or MBP-OSGIN1-6xHis. The right lanes show 20% input for each sample.

We employed an mNeonGreen (mNG)-based bimolecular fluorescence complementation (BiFC) approach to further ascertain these results and assess whether dimeric interactions can occur in cells. This approach relies on the fusion of two non-fluorescent mNG fragments (domains 1-10 and 11) to proteins that, upon interacting, allow the reconstitution of mNG fluorescence (Feng et al., 2017). Using this approach, we measured significantly higher fluorescence levels in *OSGIN1*-deleted HeLa cells that co-expressed OSGIN1-mNG^1-10^ with OSGIN1-mNG^11^ compared to mNG^11^ alone (Figure 7A-C). Similar results were obtained in *OSGIN2*-deleted HeLa cells co-expressing OSGIN2-mNG^1-10^ with OSGIN2-mNG^11^ (Figure 7A-C). Furthermore, we found high fluorescence levels in *OSGIN1/2*-deleted HeLa cells co-expressing OSGIN1-mNG^1-10^ with OSGIN2-mNG^11^ as compared to control (Figure 7A-C). Interestingly, we found that both homo- and heteromers containing OSGIN2 showed an increase of fluorescence signal at the intercellular bridge in late mitosis (Figure 7D-E). These results support those obtained *in vitro* and indicate that OSGIN1 and OSGIN2 can form both homo- and heteromeric complexes in cells. They further suggest that some of these complexes can localize to the intercellular bridge in late mitosis, supporting the notion that they locally exert their function to impact its stability.

**Figure 7.**
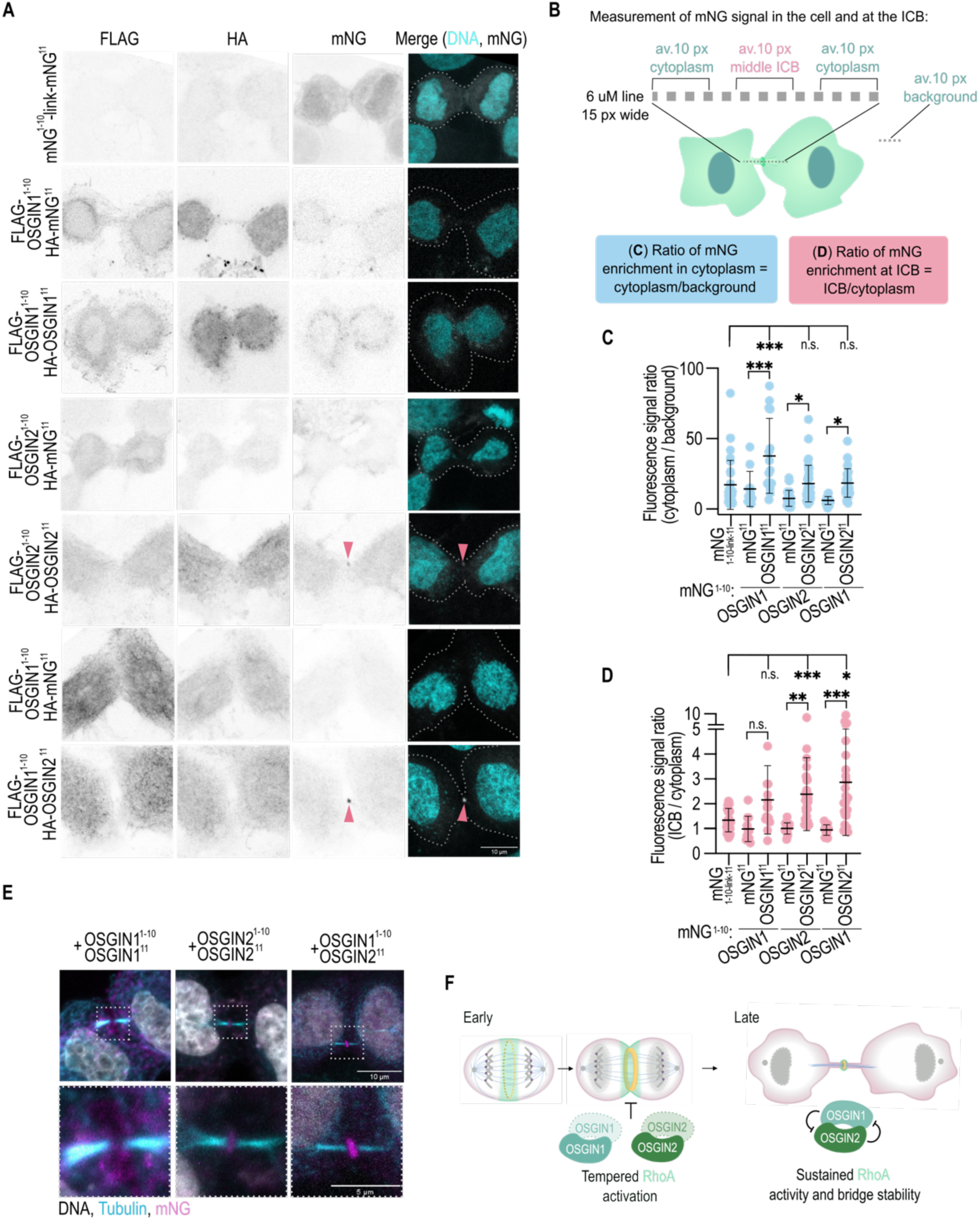
OSGIN1 and OSGIN2 can form homo- and heteromeric complexes in cells. (A) Representative confocal images of fixed control HeLa cells in telophase expressing mNG^1-10^-link-mNG^11^ (positive control), *OSGIN1(-)* cells co-expressing mNG^1-10^-3xFLAG-OSGIN1 with either HA-mNG^11^ or HA-OSGIN1-mNG^11^, *OSGIN2(-)* cells co-expressing mNG^1-10^-3xFLAG-OSGIN2 with either HA-mNG2^11^ or HA-OSGIN2-mNG^11^, and *OSGIN1/2(-)* cells co-expressing mNG^1-10^-3xFLAG-OSGIN1 with either HA-mNG^11^ or HA-OSGIN2-mNG^11^. Cells were stained with anti-FLAG, anti-HA and Hoechst 33342 (revealing nuclei in the merged images, cyan). The mNG fluorescence signal shown is native. Arrowheads point to fluorescence signal at the intercellular bridge. Scale bar = 10 µm. (B) Schematic depiction of the quantification method employed to measure fluorescence signal in the cytoplasm (C) and at the intercellular bridge (D). (C-D) Quantification of the fluorescence signal ratio of cytoplasm over background (C) or intercellular bridge over cytoplasm (D) in the conditions shown in (A). n=11-37 cells acquired in N ≥ 3 replicates. n.s. = not significant. \**p* < 0.05, \*\**p* < 0.01, \*\*\**p* < 0.001, one-way ANOVA with Sidak’s multiple comparison correction. (E) Representative indirect immunofluorescence images of fixed HeLa cells in telophase co-expressing either mNG^1-10^-3xFLAG-OSGIN1 with HA-OSGIN1-mNG^11^, mNG^1-10^-3xFLAG-OSGIN2 with HA-OSGIN2-mNG^11^, or mNG^1-10^-3xFLAG-OSGIN1 with HA-OSGIN2-mNG^11^. Cells were stained with anti-α-tubulin (cyan) and Hoechst 33342 (revealing nuclei, white). The mNG fluorescence signal shown (magenta) is native. The boxed region is magnified 3 time on the bottom panels. Scale bar = 10 µm (5 µm in magnified panel). (F) Schematic illustration of the proposed model for OSGIN protein function during cytokinesis. OSGIN1 and OSGIN2 redundantly limit RhoA activity in early cytokinesis and antagonize one another in late cytokinesis, sustaining intercellular bridge stability.

## Discussion

Here, we characterize the human OSGIN1 paralog OSGIN2, revealing it is a novel FMO required for cytokinesis completion. We find that cells lacking either OSGIN1 or OSGIN2 show intercellular bridge instability and decreased RhoA activity in late cytokinesis. Initially this suggested non-redundant functions for the two genes, but we found that multinucleation defects are suppressed in cells mutant for both *OSGIN1* and *OSGIN2*, indicating that the two genes partially antagonize each other. Consistent with this, we found that both OSGIN1 and OSGIN2 are capable of homo- and heteromerization and that their interaction is likely direct, suggesting they regulate each other’s function. Furthermore, we show that *OSGIN1/2*-depleted cells have increased RhoA activity at the cell equator in early cytokinesis, indicating that both OSGIN1 and OSGIN2 downregulate the activity of the small GTPase RhoA. Taken together, our results support a model (Figure 7F) in which OSGIN1 and OSGIN2 both act as negative regulators of RhoA signaling in early cytokinesis, and that their heteromeric interaction antagonizes this activity in late cytokinesis to favor intercellular bridge stability and cell abscission.

Our model posits that OSGIN1 and OSGIN2 redundantly limit RhoA activity in early cytokinesis, which is supported by our finding that the equatorial levels of the RhoA activity sensor are significantly increased at the onset of cytokinetic furrow ingression in *OSGIN1/2*(-) cells. We found that furrow ingression occurs sooner in these double mutant cells compared to control cells, which is likely consequential to the increase in RhoA activity (David et al., 2014). *OSGIN1/2*(-) cells also have higher levels of RhoA activity at the intercellular bridge in late cytokinesis compared to *OSGIN1*(-) and *OSGIN2*(-) single mutant cells and undergo abscission sooner than control cells. While an overall acceleration of cytokinesis progress could account for the phenotypic suppression that we observe in *OSGIN1/2*(-) cells, we consider this possibility as unlikely because *OSGIN1*(-) cells undergo furrow regression even sooner than the measured time of abscission in *OSGIN1/2*(-) cells. We rather favor a view in which the mutual antagonistic regulation of OSGIN1 and OSGIN2 in late cytokinesis is key to prevent the inappropriate downregulation of RhoA activity at this stage, thus sustaining intercellular bridge stability. Our finding that OSGIN1 and OSGIN2 heteromers accumulate at the intercellular bridge in late cytokinesis further support this view.

One key aspect of the antagonism between OSGIN1 and OSGIN2 is that it manifests itself in late cytokinesis. This suggests that OSGIN1 and OSGIN2 heteromer formation becomes more significant as cells progress through mitosis and culminates in late cytokinesis. The mNG-based BiFC approach that we employed to characterize oligomers *in vivo* does not enable us to resolve this because the mNG fragments that reconstitute fluorescence remain stably bound to one another once they are associated (Feng et al., 2019; Koker et al., 2018), thus precluding temporal analysis. Mitotic exit is characterized by changes in several cellular regulators, including mitotic kinases and regulators of the abscission checkpoint, that can differentially impact the outcome of cytokinesis (Gibieza and Petrikaite, 2024). While it is unknown whether one or both OSGIN proteins respond to such mitotic regulators, their accumulation at the intercellular bridge in late cytokinesis suggests that they respond to cell cycle-dependent activities.

Our findings show that OSGIN proteins form both homo- and heteromeric complexes and, at least for OSGIN1, that this interaction is direct. Genetic analysis indicates that the heteromerization of OSGIN1 with OSGIN2 results in mutual antagonism, however it is unclear whether individual OSGIN proteins function as monomers or homodimers to limit RhoA activity. While FMOs can function as monomers, many have been reported to form dimers in order to carry out their activity (Bailleul et al., 2023; Cho et al., 2011; Kachalova et al., 2010; Nicoll et al., 2020; Richardson et al., 2024; Schreuder et al., 1988). The identification of domains and residues mediating interactions in OSGIN proteins will be key to disentangle the relative contribution of oligomerization in OSGIN protein activity and function.

Vertebrates have two distinct genes encoding OSGIN proteins, while most invertebrates have a single one. In *C. elegans*, where the role in cytokinesis was uncovered, loss of OSGN-1, the OSGIN1 ortholog and sole OSGIN protein, results in a decrease of RhoA activity at the stable bridge formed between the two nascent primordial germ cells (Goupil et al., 2017; Goupil et al., 2024), a phenotype opposite to what we find after deleting both *OSGIN* genes in human cells. The OSGIN loci in both organisms are predicted to transcribe alternative mRNA species, and thus it is possible that *C. elegans* regulates the activity of OSGN-1 via an alternative, antagonistic form of the protein that dimerizes with the canonical one. Alternatively, while cytokinesis itself is highly conserved, it involves several regulators whose requirements in the process have been shown to vary between cell types and species (Husser et al., 2022; Ozugergin et al., 2022; reviewed in Ozugergin and Piekny, 2022). Another possibility is therefore that duplication of the *OSGIN* locus in vertebrates afforded the adaptation of OSGIN protein activity to specific demands or conditions.

Finally, the expression of both OSGIN genes was shown to be elevated under oxidative stress, and while little is known about the cellular role of OSGIN2 outside of cytokinesis, OSGIN1 has been implicated in apoptosis, ferroptosis and autophagy in certain conditions (Deng et al., 2025; Hu et al., 2012; Jia et al., 2024; Sukkar and Harris, 2017; Tang et al., 2024; Wang et al., 2017; Yan et al., 2026; Yao et al., 2008; reviewed in Hussey et al., 2025; Kim et al., 2025). It will be interesting to assess whether these activities depend on the monooxygenase activity of OSGIN proteins or are impacted by OSGIN protein oligomerization, to determine whether these properties are general features of OSGIN protein function or have been favored in the context of cytokinetic regulation.

## Materials and Methods

### DNA constructs and cloning

The vectors and oligonucleotides used in this study are available in the Tables S1 and S2. The constructs detailed below were achieved through PCR amplifications using Phusion DNA polymerase (NEB, M0530L), following the manufacturer’s instructions, and sequences were validated by Sanger sequencing.

To generate the pEGFP-HA-OSGIN1-P^26^L, pEGFP-HA-OSGIN2-P^70^L and pcDNA3.1(+)-HA-HsOSGIN1-P^26^L constructs to verify their function and/or localization in cells, mutation of a single base pair enabling the desired amino acid change was realized using the NEB Q5 Site-directed mutagenesis kit (NEB, E0554S). Forward and reverse oligonucleotides containing the desired mutation were designed using the NEBaseChanger tool. Sequences were amplified by PCR from source vectors (Table SI) and then ligated following the manufacturer’s instructions.

To generate pcDNA3.1(+)zeo-TagBFP-3xFLAG-OSGIN1, pcDNA3.1(+)zeo-TagBFP-3xFLAG-OSGIN2, pCMV-iRFP670-HA-OSGIN2, pRK5-HA-OSGIN2-P^70^L, pEGFP-HA-OSGIN2S, pET-22-MBP-OSGIN1-His_6_ and the vectors for Split-mNG2 expression, inserts were amplified by PCR from sources vectors and inserted by Gibson assembly into backbone vectors amplified by PCR.

To generate lentiCRISPRv2-Hygro-O2sg4, plentiCRISPRv2-Hygro (a gift from Brett Stringer; Addgene plasmid # 98291 (Stringer et al., 2019)), was digested with BsmBIv2 (NEB R0739S) and ligated with OSGIN2 sg2 phosphorylated with T4 Polynucleotide kinase (NEB, M0201S), and annealed with duplexed oligos (Table S1), following the method developed in (Sanjana et al., 2014; Shalem et al., 2014).

### Human cell culture, transfection and lentiviral transduction

Human cell lines were grown at 37 °C with 5% CO_2_ in Dulbecco’s Modified Eagle’s Medium (Wisent, 319-005-CS) complemented with 10% (v/v) heat-inactivated fetal bovine serum (Wisent, 090150), 100 U/mL penicillin and 100 μg/mL streptomycin (Wisent, 450-201-EL).

For transient transfections, cells were seeded at a density of 5 × 10^5^ cells per 10-cm dish, 1.5-3 × 10^5^ cells per well in a 6-well plate or 35-mm microscopy plate, and transfected using Lipofectamine 3000 treatment (Invitrogen, L3000015) according to the manufacturer’s instructions. Experiments were performed 36 to 48 h post-transfection. Stable cell lines expressing GFP-HA alone, GFP-HA-OSGIN1 (WT or P^26^L) or GFP-HA-OSGIN2 (WT or P^70^L) were generated by transfection (as above) and subsequent selection for 14 days in growth medium supplemented with 1.5 mg/mL G418 sulphate solution (Wisent, 450-130-QL).

For RNAi experiments, 1 × 10^5^ HEK 293 or HeLa cells were seeded in 6-well plates, with or without a glass coverslip at the bottom. A pool of 4 siRNAs (siGENOME Human *OSGIN1* (29948) siRNA-SMARTpool, M-010197-01-0010) directed against *OSGIN1* was transfected using the Dharmacon transfection reagent DharmaFECT 1 (T-2001-02), following manufacturer’s instructions. Controls included transfection reagent alone, as well as transfection of a pool of non-targeting siRNA from manufacturer (Dharmacon, siGENOME non-targeting siRNA Pool #1, D-001206-13-05). Transfection reagent was added to a tube containing 200 μL of OPTI-MEM (Gibco, 51985034). siRNAs were mixed separately in a tube containing 0,2 mL/condition of OPTI-MEM at pre-determined concentrations (25 nM for control siRNA in all cell types, 50 nM *OSGIN1* siRNA in HEK 293 cells and 75 nM *OSGIN1* siRNA in HeLa cells). Finally, 0,4 mL of the transfection mix was added to the wells for a total volume of 2 mL per well. Cells seeded on coverslips were fixed and immunostained for multinucleation assessment. Cells seeded without a coverslip were processed for total RNA extraction for qPCR.

HeLa cells stably expressing the dTom-2xrGBD RhoA activity sensor (Mahlandt et al., 2021) were generated by lentiviral transduction using the pLV-dimericTomato-2x-rGBD vector (a gift from Dorus Gadella; Addgene #176098) in HEK 293T cells. After transfection, supernatants were treated as described above (harvested, spun, filtered, mixed with hexadimethrine bromide) and added to *OSGIN2*(-) or *OSGIN1/OSGIN2*(-) HeLa cells (*ROSA26*(-) and *OSGIN1*(-) stably expressing dTom-2xrGBD were previously generated; (Goupil et al., 2024)). After selection with 1 µg/mL puromycin for 48 h, cells positive for dTom-2xrGBD were sorted with a BD FACSAria™ cell sorter. Cells with minimal dTom-2xrGBD expression were recovered and grown as a polyclonal population.

### Editing of the *OSGIN2* locus

The deletion of *OSGIN2* in *OSGIN1*(-) HeLa cells and HEK 293 cells was done by CRISPR/Cas9-mediated gene editing, as described previously (Goupil et al., 2024; Ran et al., 2013; Sanjana et al., 2014). A 10-cm dish was first seeded with 1.5 × 10^6^ HEK 293T cells, which were grown for 24h. The following day, cells were co-transfected (using Lipofectamine 3000 treatment, as described above) with pMD2.g (envelope plasmid, Addgene #12259), psPAX2 (packaging plasmid, Addgene #12260, a gift from Didier Trono), and lentiCRISPRv2-Hygro or lentiCRISPRv2-puro (transfer plasmid) containing a small guide (sg) RNA for human OSGIN2 (sg4). The next day, the medium was changed to 20% FBS-containing medium and the cells were incubated for an additional 36h. The medium was then harvested, centrifuged for 5 min at 860 g to remove debris, filtered with a 0.45 µm surfactant-free cellulose acetate syringe filter (Corning, 431220) and supplemented with hexadimethrine bromide (Sigma 107689) to a final concentration of 8 µg/mL. To transduce cells, the filtered lentivirus-containing medium was applied directly to either *OSGIN1*(-) HeLa cells or HEK 293 cells seeded 24 h prior at a density of 1 x 10^6^ cells in a 10-cm dish. The next day, the medium was changed to medium containing 1 µg/mL puromycin (Wisent 400-160-EM) only (for *OSGIN2*(-) HEK 293 cells) or both puromycin and 200 µg/mL hygromycin (Wisent 400-141-XL) (for *OSGIN1/OSGIN2*(-) HeLa cells) for selection of transduced cells.

The *OSGIN2* editing efficiency was verified by extracting genomic DNA (Monarch Genomic DNA Purification Kit, NEB T3010S) and amplifying by PCR the region of the gene targeted by the sgRNA. Cells were then singled out in 96-well plates in conditioned medium containing 20% FBS (using a BD FACSAria™ cell sorter), and individual clones were allowed to expand before genomic DNA was extracted and PCR-amplified again to sequence part of exon 4 of *OSGIN1* and exon 2 of *OSGIN2* (see oligonucleotide sequences in Table 2) and analyzed with Synthego’s ICE tool. For *OSGIN1/OSGIN2* (-) HeLa cells, one recovered clonal population with >90% of cells bearing a deletion in *OSGIN1* and *OSGIN2* was considered as being mutant for both genes and subjected to all analyses. HEK 293 cells deleted of *OSGIN2* were screened following a similar approach, from a polyclonal population with 86% of cells bearing a genomic deletion in exon 2. After singling out in 96-well plates, a monoclonal population was recovered and shown to have 90% of cells bearing a deletion in *OSGIN2*.

### RNA isolation and quantitative PCR analysis

HeLa or HEK 293 cells were plated in 6-well plates as described above, grown at 37°C and treated for RNAi depletion for 48h. Total RNA extraction was performed using the RNeasy Mini Kit (Quiagen, 74104) following manufacturer’s instructions. 2 µg of total RNA was used for the synthesis of cDNA, which was done using the High-Capacity cDNA Reverse Transcription kit (Applied Biosystems, 4368814). Two technical replicates of qPCR were performed with 20 ng of cDNA per reaction using the Taqman Fast qPCR MasterMix (Applied Biosystems, 4444557). Reactions were performed for 40 cycles on a QuantStudio™ 7 Flex Real-Time PCR System using an *OSGIN1*-specific primer set from the Universal ProbeLibrary (see Table II). Data were analyzed with Expression Suite software (Applied Biosystems), using ACTB and GAPDH as reference genes.

### Indirect immunofluorescence

HeLa or HEK 293 cells were grown as asynchronous populations in 6-well plates with glass coverslips (1-2 x 10^5^ cells/well) for 24 h prior to transient transfection (using Lipofectamine 3000 treatment, see above). Medium was aspirated 48 h later and cells were fixed for 5 min at RT with paraformaldehyde (4% v/v in PBS buffer, 137 mM NaCl, 2.7 mM KCl, 8 mM Na_2_HPO_4_, and 2 mM KH_2_PO_4_). Fixed cells were washed three times in PBS buffer, permeabilized for 5 min at RT in 0.3 % Triton X-100 in PBS buffer, and blocked for 20 min at room temperature (RT) in PBS buffer supplemented with 2% w/v BSA. The cells were then incubated for 1h at RT with primary antibodies diluted in PBS buffer supplemented with 0.5% BSA, washed 3 times with PBS buffer, and incubated for 1h at RT with Alexa fluor dye-conjugated secondary antibodies (1:500) and Alexa-555-coupled Phalloidin (1:400, Invitrogen, A34055) (for experiments assessing binucleation levels), diluted in PBS buffer supplemented with 0.5% BSA. After three washes with PBS buffer, cells were treated for 5 min at RT with Hoechst 33342 (1 µg/mL in PBS buffer, Invitrogen, H1399), followed by two additional washes of 10 min each with PBS buffer. For split mNeonGreen (mNG) experiments, cells were stained against HA and FLAG to validate constructs expression, and the endogenous mNG signal was not amplified with antibodies. Coverslips were mounted/sealed on glass slides using Mowiol mounting agent [200 mM Tris pH 8.5, 9.8% w/v of Mowiol (Sigma, 81381), 24% w/v glycerol, 0.02% w/v sodium azide] and allowed to dry overnight at RT before imaging. The following primary antibodies were used: goat α-GFP (1:1000, Rockland Immunochemicals, 600-101-215) rat α-HA (1:100, Sigma, 11867423001), rabbit α-pMLC-S20 (1:50, Abcam, AB2480), mouse α-α-tubulin (clone DM1A, 1:5,000, Sigma, T6199), mouse α-FLAG M2 (1:400, Sigma, F3165). The following Alexa Fluor-coupled secondary antibodies were used: donkey α-rabbit 488-conjugated (Invitrogen, A21206), donkey α-mouse 647-conjugated (Invitrogen, A31571), donkey α-goat 488-conjugated (Invitrogen, 11055), donkey α-mouse 546-conjugated (Invitrogen, A10036), donkey α-rat Cy3-conjugated (Jackson Immunoresearch, 712-165-153) and donkey α-rat Cy5-conjugated (Jackson Immunoresearch, 712-175-150).

### Protein expression, purification and pull-down in vitro

*Escherichia coli* BL21 (DE3) bacteria transformed with either pET-22-MBP-OSGIN1-His_6_ or pET-22-OSGIN1-His_6_ were grown in LB broth at 37 °C until an O.D. of ∼0.7 to 0.8 was reached, and protein production was induced with 0.5 mM IPTG with a supplement of 0.05 mM riboflavin (Sigma, R9504) overnight at 18 °C. All subsequent steps were performed on ice. Cells were collected by centrifugation, resuspended in lysis buffer (50 mM Tris pH 8.0, 150 mM NaCl, 20 mM imidazole, 1 mM DTT, 1 mM PMSF, 0.1% Triton X-100) using 1/10 of the volume of the culture medium and sonicated 6 x 10 s (at 50% intensity, using a Fisher Scientific FB120 sonicator). To remove cell debris, lysates were spun at 18,000 × g, and then incubated with Nickel-NTA agarose (QIAGEN, 1018244) for 2h at 4 °C. Beads were washed thrice in 10 bed volumes of high salt buffer (20m M Tris pH 8.0, 500 mM NaCl, 5 mM B-ME, 20 mM imidazole, 5 mM MgCl2) and once in ten bed volumes of low salt buffer (20m M Tris pH 8.0, 150 mM NaCl, 5 mM β-ME, 20 mM imidazole, 5 mM MgCl2). Bound material was eluted from the agarose matrix by incubating in 7 mL elution buffer (20 mM Tris pH 8.0, 150 mM NaCl, 5 mM β -ME, and 300 mM imidazole), thrice 5 min at 4 °C. Eluates were concentrated using a Sartorius Vivaspin 20 Centrifugal Concentrator (10k MWCO, Fisher, 14-558-501) and passed through a Zeba spin desalting column (7k MWCO, Thermo Scientific, PI89892) saturated with storage buffer (20 mM Tris pH 7.5, 150 mM NaCl, 1mM DTT, 10% v/v glycerol), dosed using a Bradford assay, and finally aliquoted and flash frozen in liquid nitrogen for further use. Protein purity was verified by SDS-PAGE, followed by Coomassie staining.

For pull down assays, all steps were performed on ice. Equimolar amounts of purified MBP-His_6_ or MBP-OSGIN1-His_6_ were incubated in 0.5 mL of pull-down buffer (20 mM HEPES, 10 mM imidazole, 1 mM DTT) supplemented with 0.5% Triton X-100 and protease inhibitors (phenylmethanesulfonyl fluoride (PMS444), Aprotinin (APR200), Leupeptin (LEU001), Pepstatin A (PEP605), all from Bioshop Canada) added fresh, with amylose resin (NEB, E8021S) (which had been previously washed thrice in pull-down buffer) on a rotating wheel for 2h at 4°C. The resin and bound proteins where then spun 2 min at 4000 rpm, the supernatant was discarded and the resin was resuspended in pull-down buffer supplemented with 3% w/v BSA and incubated on a rotating wheel for 30 min at 4°C for blocking. The resin and bound proteins were spun again (4000 rpm – 2 min) and washed thrice with 0.5 mL of pull-down buffer. Input sample representing 20% of the total volume of resin were taken and resuspended in Laemmli 2X buffer. The resin and bound proteins were then incubated with an equimolar amount of OSGIN1-His_6_ (2 uM final for each) in pull-down buffer (diluted from a 10X preparation to maintain concentrations in final volumes) and incubated on a rotating wheel for 2h at 4°C. The resin and bound proteins were spun (4000 rpm – 2 min) and washed thrice with 0.5 mL of pull-down buffer and resuspended in Laemmli 2X buffer and visualized via SDS-PAGE and Coomassie staining.

### Immunoprecipitation

For co-immunoprecipitation experiments, HEK 293 cells were seeded in a 10 cm dish at a density of 5 × 10^5^ cells (2 dishes per condition). Cells were transfected the next day using polyethylenimine (PEI, Sigma, 764965, at a 1 DNA:3 PEI ratio). Experiments were performed 48 h post-transfection. To synchronize cells in prometaphase, media was changed 24h post-transfection for media containing 2 uM S-trityl-L-cystein (STC, Sigma, 164739). The next day, cells were washed three times with media to release from prometaphase. Washes were kept, spun (300 x g, 2 min) and pellets were also washed to remove STC, and redistributed into the plates. Cells were incubated for 90-120 min after release to let them enter telophase.

Telophase-enriched HEK 293 cells were put on ice, washed twice with cold PBS buffer, collected using a scraper and resuspended in HEPES buffer (50 mM HEPES, pH 7.4, 50 mM NaCl, 5 mM EDTA, 10% v/v glycerol) supplemented with 1% v/v Triton-X-100 and protease inhibitors freshly added. Detached cells present in the supernatants from plate washes were also collected, washed and added to the scraped samples. The samples were incubated for 20 min at 4°C on a rotating wheel for lysis, and clarified using a tabletop centrifuge (16 x g) for 20 min at 4°C. Ten percent of the supernatant was kept for SDS-PAGE analyses (total cell lysate) and added to Laemmli 5X buffer (250 mM Tris pH 6.8, 10% SDS, 30% (w/v) glycerol, 0.01% bromophenol blue) and the rest was incubated with protein G-agarose (BioShop, PRA286.5) on a rotating wheel for 30 min at 4°C to remove non-specific binding (pre-clearing). Then, lysates were spun (4000 rpm – 2 min), and supernatants were incubated with mouse α-FLAG M2 (1/800, Sigma, F3165) and protein G-agarose on a rotating wheel overnight at 4°C. The next day, beads were washed three times in HEPES buffer and Laemmli 2X buffer was added to elute the immunoprecipitated material prior to SDS-PAGE.

### Gel electrophoresis and immunoblotting

Protein samples were run on 10% SDS-PAGE and transferred to nitrocellulose. Membranes were blocked for 1h at RT in TBST buffer (150 mM NaCl, 50 mM Tris pH 7.6, 0.1% Tween 20) supplemented with 5% powder milk, followed by blotting overnight at 4°C with primary antibodies in TBST buffer. Membranes were washed 3 x 10 min in TBST buffer, incubated at RT with horseradish peroxidase-coupled secondary antibodies for 1h in TBST buffer supplemented with 5% powder milk, and washed again 3 x 10 min in TBST buffer. Samples were revealed using Clarity Western ECL Substrate (Biorad, 1705060) and ImageQuant LAS 4000 Imager (GE Healthcare). The following primary antibodies were used: mouse α-α-tubulin (clone DM1A, 1:5,000, Sigma, T6199), rat α-HA (1:1000, Sigma,11867423001), mouse α-FLAG M2 (1:4000). The following secondary antibodies were used: goat α-mouse-HRP (1:10,000, Bio-Rad, 170-6515) and goat α-rat-HRP (1: 5000, Sigma, AP136P).

### Microscopy and live-cell imaging

For fixed cells, images were acquired with a HC PL APO CS2 63×/1.4 NA oil-immersion objective mounted on a Leica SP8 laser-scanning confocal microscope, and 0.3 to 0.8-µm-thick confocal sections of the entire cells obtained by illumination in multi-track mode with 405, 488-, 552 and 638-nm laser lines controlled by LAS X software (Leica).

For live-cell imaging, HeLa cells were plated on 35-mm microscopy dishes (MatTek, P35G-1.5-14-c) and left to adhere overnight before co-transfection (using Lipofectamine 3000 treatment, as above) of vectors enabling expression of mRFP-CAAX, iRFP670 alone or iRFP670-HA-OSGIN2, or Citrine-αTubulin. For dimericTomato-rGBD imaging or *ROSA26*(-), *OSGIN1*(-), *OSGIN2*(-) or *OSGIN1/2*(-) bright field imaging, which required no transient transfection, cells were plated and grown for 24-48 h before imaging.

For furrow ingression measurements, cells in prometaphase or metaphase were selected from asynchronous populations 48h after seeding. Time-lapse images were acquired for 10h at 5-min intervals with a Plan-Apocromat 40x/1.4 NA oil-immersion objective mounted on a Zeiss LSM700 laser-scanning confocal microscope controlled by Zen software, acquiring 0.8-µm-thick confocal sections of the entire cell using single-track mode illumination with 555-nm laser line.

For measurements of membrane stability and membrane scission, time-lapse images were acquired for 18h at 10-min intervals with a HC PL APO CS2 20x/0.75 DRY objective mounted on a Leica SP8 laser-scanning confocal microscope controlled by LAS X software (Leica), acquiring 0.8 µm-thick confocal sections using transmitted bright-field illumination.

For measurements of midbody enrichment of iRFP670 alone, iRFP670-HA-OSGIN2, or the dTom-2xrGBD reporter or the measurement of midbody microtubules clearing with Citrine-αTubulin, time-lapse images were acquired for 10h at 3-min intervals with a HC PL APO 63X/1.4 N.A oil-immersion objective mounted on a Leica SP8 laser-scanning confocal microscope controlled by LAS X software (Leica), acquiring 0.8-µm-thick confocal sections of the entire cell using single-track mode illumination with 488-, 552-nm or 638-nm laser lines. Anaphase onset (t = 0 min) was again defined as the first frame where two sets of chromosomes are visible or the frame where the cell shape transitions from round to elongated.

### Image analysis and fluorescence quantification

All images were analyzed and processed for figures from original files using ImageJ software (NIH). Analyses were performed on sum or maximal intensity projections converted into black and white renderings. To improve visualization in chosen figures brightness and contrast were adjusted using the same settings for all images within a panel.

Multinucleation analysis was performed on TIFF images from microscopy files using ImageJ Cell Counter plugin. For rescue experiments with GFP constructs, only cells with cytoplasmic fluorescence signal in the 488 channel were included in the count.

Cytokinetic furrow ingression was measured by tracking plasma membrane signal in HeLa cells (mRFP-CAAX probe) on a single z-slice at the center of the cell, for every time point until 60 minutes after anaphase onset. A 3 pixel-wide line was traced through the cell’s equator, and the distance (in microns) between the fluorescence peaks representing membranes was measured at each time point.

mNeonGreen2 fluorescence analysis was performed by tracing a 0.75 µm thick and 6 µm-long line across the intercellular bridge on a max projection of 4 z-slices. The average of fluorescence gray value of 0.5 µm at each extremity of this line was used as measurement for cytoplasmic fluorescence. The average fluorescence gray value of 0.5 µm at the middle of the line (corresponding to the middle of the intercellular bridge) was used as measurement for intercellular bridge enrichment. A 0.75 µm-thick line was traced in the background (outside of the cytoplasm) and the average fluorescence gray value along this line was used as a background value. To assess cytoplasmic enrichment of mNeonGreen fluorescence, a ratio between the cytoplasmic fluorescence value and the acquired background value was calculated. To assess intercellular bridge enrichment of mNeonGreen fluorescence, a ratio between the intercellular bridge fluorescence value and the cytoplasmic value was calculated.

For measurements of dTom-2xrGBD sensor levels in dividing HeLa cells at the furrow and intercellular bridge, a 0.25 µm-thick line was drawn across the cell equator on sum projections and fluorescence integrated density was acquired for every timepoint from 3 minutes before anaphase onset up to 30 minutes and every 5 timepoints between 30 and 60 minutes. For the same timepoints, integrated density was also measured in the cytoplasm of each daughter cell along 2 lines drawn on each side of the equator. Sensor enrichment was quantified as the ratio between the value for peak fluorescence intensity at the equatorial cortex and the mean value found at the cortical regions on either side within each daughter cell.

For measurements of dTom-2xrGBD sensor distribution at the equator, fluorescence gray value was measured along a 15 µm-wide line which was drawn horizontally across the dividing cell from the cytoplasm of one daughter cell to the other. Cytoplasmic background was measured with an average of the three lowest cytoplasmic values obtained. The background value was deduced from highest peak value (the average of 2 µm surrounding the highest peak), and 50% of this obtained value (peak minus background) was used as a threshold to score the distance over which the signal spreads at the equator. The length (uM) along the line within the established threshold (where values were 50% or higher of the peak value) was recorded.

For measurements of iRFP760 enrichment at the equator and intercellular bridge, a horizontal line was traced across the dividing cell on a maximum projection, for selected time points. Equatorial fluorescence value was obtained by averaging 3.75 µm at the line midpoint. Cytoplasmic fluorescence values were obtained by averaging 3.75 µm at each extremity of the line. Equatorial enrichment was determined as the ratio between the fluorescence value at the equator and the value in the cytoplasm.

For cytokinetic outcome and membrane scission analysis, the first time point after cytokinetic ingression (i.e when the bridge is fully ingressed) was considered T=0. Membrane regression timing was defined as the first frame where the partition between the dividing cells was ruptured. Membrane scission timing was defined by the first frame where the intercellular bridge was no longer continuous from one cell to the other and/or no longer visible.

### Statistical analysis

Graph design and all statistical analyses were realized with the GraphPad Prism software. Parametric tests of statistical significance between samples were applied except when assumptions of normality and equal variance were not met; see figure legends for details. In all cases, a two-tailed *p*-value smaller than 0.05 was considered significant. All results are expressed as average ± SD or SEM. Sample size (n) for each experiment is indicated in each figure panel or legend. At least three independent biological replicates (N) were realized for each condition in all experiments.

## Acknowledgments

We thank Drs Michel Bouvier and Gregory Emery for constructs, cell lines and reagents, and Maxime Roussel for assistance with immunoprecipitations. We also thank Christian Charbonneau, Annie Gosselin, Angélique Bellemare-Pelletier and Raphaëlle Lambert of the Institute for Research in Immunology and Cancer (IRIC) Bio-imaging, Flow cytometry and Genomics Facilities for technical assistance, and members of the Smith and Labbé laboratories for helpful discussions. L.L. was a recipient of studentships from the Fonds de la Recherche du Québec - Santé (FRQ-S), the Canadian Institutes of Health Research (CIHR), Université de Montréal’s Graduate Studies and the IRIC. M.J.S. holds the Canada Research Chair in Cancer Signaling and Structural Biology. This study was supported by grants from the CIHR (PJT-480641) to M.J.S. and J.-C.L.

