## Supplemental information for "The flavin-containing monooxygenases OSGIN1 and OSGIN2 antagonize each other to favor cytokinesis completion"

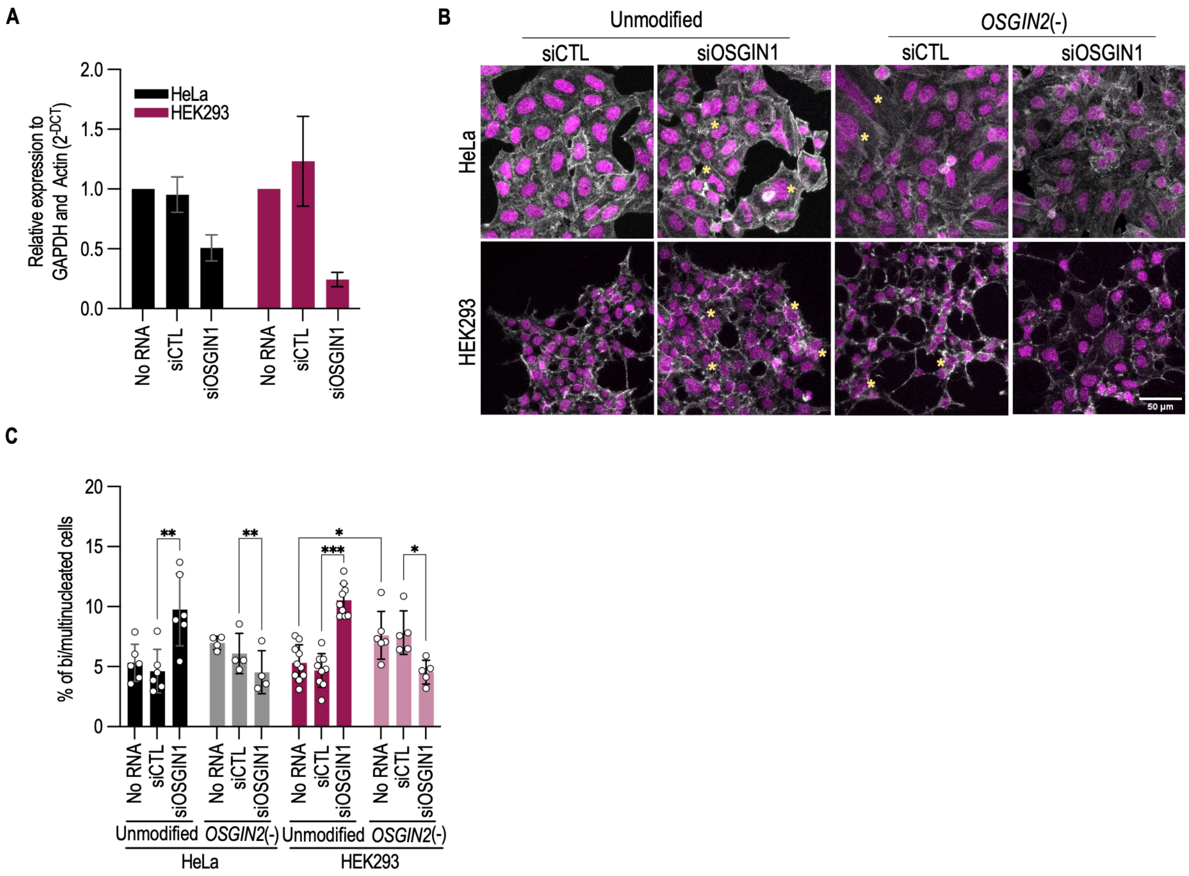


**Figure S1. OSGIN1 and OSGIN2 impact cytokinesis completion in HEK 293 cells.** (A) Quantitative PCR analysis of relative *OSGIN1* mRNA levels in HeLa and HEK 293 cells following treatment with pools of 4 different siRNAs, either scrambled (siCTL) or targeting *OSGIN1*. In each cell line, *OSGIN1* levels are normalized to those of GAPDH and actin in untreated samples. Bars denote SD in N=3 replicates. (B) Representative immunofluorescence images of unmodified or *OSGIN2*(-) HeLa and HEK 293 cells, with pools of 4 different siRNAs, either scrambled (siCTL) or targeting *OSGIN1*. Cells were labelled with phalloidin (revealing F-actin, white) and Hoechst 33342 (revealing nuclei, magenta). Asterisks denote bi/multinucleated cells. Scale bar = 50 µm. (C) Quantification of the proportion of bi/multinucleated in unmodified or *OSGIN2*(-) HeLa and HEK 293 cells, either untreated (no RNAi) or treated with scrambled (siCTL) or OSGIN1 siRNA, as indicated. n=124-872 cells scored in N ≥ 3 replicates. Bars denote SD. *$p < 0.05$, **$p < 0.01$, ***$p < 0.001$, one-way ANOVA with Tukey's multiple comparison correction.

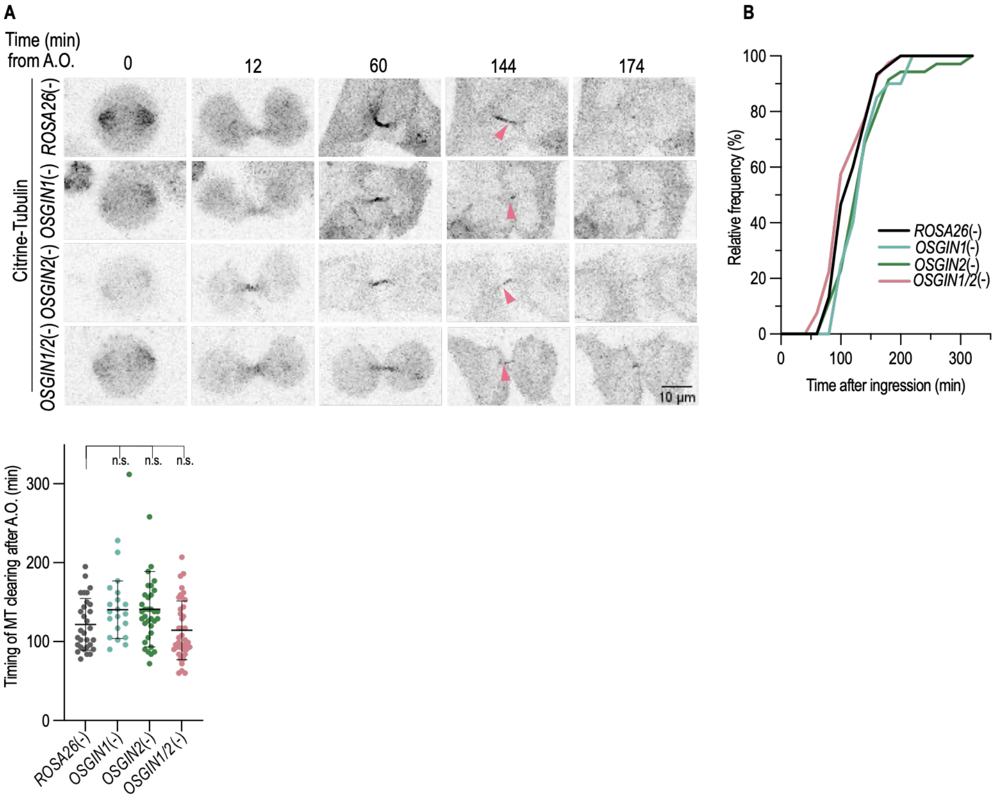


**Figure S2. OSGIN1 and OSGIN2 do not impact the timing of microtubules severing prior to abscission.** (A) Time-lapse confocal images of Citrin-tubulin fluorescence signal in dividing *ROSA26*(-) control, *OSGIN1*(-), *OSGIN2*(-) and *OSGIN1/2*(-) HeLa cells. Arrowheads indicate severed microtubule signal at the intercellular bridge. Time (in min) is relative to anaphase onset (A.O.). Scale bar = 10 µm. (B-C) Cumulative frequency of cells in which microtubules at the intercellular bridge are severed over time (B) quantification of the timing of microtubule severing relative to A.O. (C) in the same conditions as in (A). n=20-40 cells acquired in N=3 replicates. Bars denote mean ± SD. n.s. = not significant, one-way ANOVA with Dunnett's multiple comparison correction.

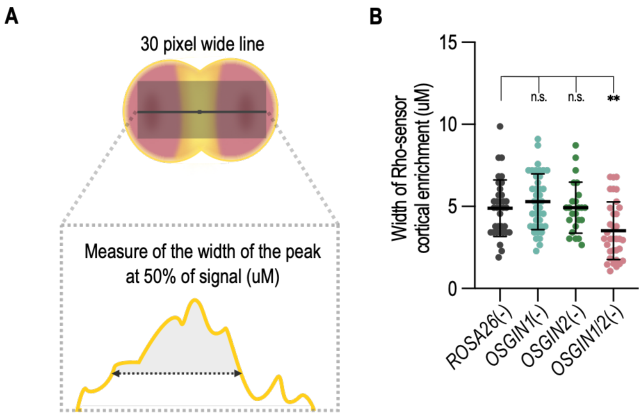


**Figure S3. OSGIN1 and OSGIN2 impact the distribution of RhoA at the equatorial cortex.** (A-B) Schematic illustration depicting the method used (A) and quantification of the width the of RhoA activity sensor fluorescence signal at the cell equator (B) acquired 6 min after anaphase onset. Bars indicate mean ± SD. n.s. = not significant, ***p* < 0.01, one-way ANOVA with Sidak's multiple comparison correction.

### Table S1. DNA constructs

| Constructs | Method / reference | Backbone source | Oligos (5'-3') for backbone amplification | Insert source | Oligos (5'-3') for insert amplification |
|---|---|---|---|---|---|
| plentiCRISPRv2-OSGIN2sg4-puro | (Goupil et al., 2024) | - | - | - | - |
| plentiCRISPRv2-OSGIN2sg4 - Hygro | (Sanjana et al., 2014; Shalem et al., 2014) | plentiCRISPRvs-Hygro | - | - | Oligo1_caccgGTCACCAAAGTATTGCACTA Oligo2_aaacTAGTGCAATACTTTGGTGACc |
| pEGFP-HA | (Goupil et al., 2024) | - | - | - | - |
| pEGFP-HA-OSGIN1 | (Goupil et al., 2024) | - | - | - | - |
| pEGFP-HA-OSGIN2 | (Goupil et al., 2024) | - | - | - | - |
| pEGFP-HA-OSGIN2S | Gibson assembly | pEGFP-HA-HsOSGIN1 | F_GGAGGAGATGGGATAGCTTAAGGTACCGCGGGCCCG<br>R_AGTTTCTTCAACTAATGGTTGAGCTCGAGATCTGAGTCC | pET-22-OSGIN2S (previously cloned in pET-22 also by Gibson) | F_CTCAGATCTCGAGCTCAACCATTAGTTGAAGAAACTTCTTTAC<br>R_CGGGCCCGCGGTACCTTAAGCTATCCCATCTCCTCC |
| pEGFP-HA-OSGIN1-P26L | Site-directed mutagenesis | pEGFP-HA-OSGIN1 | F_GTAGGCAACGGCCTTTCAGGAATCTG<br>R_CAGATTCCTGAAAGGCCGTTGCCTAC | - | - |
| pEGFP-HA-OSGIN1-P70L | Site-directed mutagenesis | pEGFP-HA-OSGIN2 | F_AGGAAATGGACTGTCAGGAATATGCC<br>R_ATTATTACCACAGGAAAAGTC | - | - |
| pCMV-iRFP670-HA-OSGIN2 | Gibson assembly | pCMV-Kan/Neo-iRFP670 (previously cloned in pCMV by Gibson, from pLenti-LifeAct-iRFP760, Addgene #84385, a gift from Gregory Emery) | F_GAGATGGGATAGCTTAAGGTCTTTAAAAAACCTCCCACACCTCCCCCTGA<br>R_TCTGGAACATCGTATGGGTAGCGTTGGTGGTGGGCGGCGG | pEGFP-HA-OSGIN2 | R_CCGCCGCCCACCACCAACGCTACCCATACGATGTTCCAGA<br>F_GTGTGGGAGGTTTTTTAAAGACCTTAAGCTATCCCATCTCC |
| pcDNA3.1(+)-HA-HsOSGIN1-P26L | Site-directed mutagenesis | pcDNA3.1(+)-HA-HsOSGIN1 (previously cloned in pCDNA3.1 also by Gibson) | F_GTAGGCAACGGCCTTTCAGGAATCTG<br>R_CAGATTCCTGAAAGGCCGTTGCCTAC | - | - |

| | | | | | |
|---|---|---|---|---|---|
| pRK5-HA-OSGIN2-$P^{70}L$ | Restriction enzyme digestion and ligation | pRK5-HA- | ClaI and SalI restriction sites digestion | pEGFP-HA-OSGIN2-P70L | F_TTCTATCGATATGTACCCATACGATGTTCCAGATTACGCTTCC<br><br>R_GCAGGTCGACTTAAGCTATCCCATCTCCTCCTCC |
| pcDNA3.1(+)-HA-HsOSGIN1 | Gibson assembly | pcDNA3.1(+)-zeocin | F_GGAAACCGCCGTAAGTCGACCGCTCGAGCATGCATCTAGA<br>R_TGGGTAcatATCGATAGAACCCGAGCTCGGTACCAAGCTT | pRK5-HA-OSGIN1 | F_AAGCTTGGTACCGAGCTCGGGTTCTATCGATATGTACCCATACGA<br>R_TCTAGATGCATGCTCGAGCGGTCGACTTACGGCGGTTTCC |
| pcDNA3.1(+)-HA-HsOSGIN2 | Gibson assembly | pcDNA3.1(+)-zeocin | F_AGTCGACCTGCAGAAGCTTGCGCTCGAGcatgcatctAGA<br>R_TGGGTAcatATCGATAGAACCCGAGCTCGGTACCAAGCTT | pRK5-HA-OSGIN2 | F_AAGCTTGGTACCGAGCTCGGGTTCTATCGATATGTACCCATACGA<br>R_TCTAGATGCATGCTCGAGCGCAAGCTTCTGCAGGTCGACT |
| pLV/CMV-dimericTomato-2x-rGBD | (Mahlandt et al., 2021) Addgene #176098 | - | - | - | - |
| pcDNA3.1(+) zeo-TagBFP-3xFlag-OSGIN1 | Gibson assembly | pcDNA3.1(+)-puro-TagBFP-2.0 (TagBFP previously cloned in pcDNA also by Gibson) | F_AGACTAGGAAACCGCCGTAATAAGCAGATATCCATCACACT<br><br>R_CCGTCGTGGTCCTTGTAGTCATTCAGTTTATGACCCAGCT | pcDNA5-FTO-TO-UltraID-3xFlag-HsOSGIN1(OSGIN1 previously cloned in pcDNA5 by Gibson) | F_AGCTGGGTCATAAACTGAATGACTACAAGGACCACGACGG<br><br>R_GTGTGATGGATATCTGCTTATTACGGCGGTTTCCTAGTCT |
| pcDNA3.1(+) zeo-TagBFP-3xFlag-OSGIN2 | Gibson assembly | pcDNA3.1(+)-puro-TagBFP-2.0 (TagBFP previously cloned in pcDNA by Gibson) | F_GAGATGGGATAGCTTAATAGGCAGATATCCATCACACTGG<br><br>R_CCGTCGTGGTCCTTGTAGTCATTCAGTTTATGACCCAGCT | pcDNA5-FTO-TO-UltraID-3xFlag-HsOSGIN2 (OSGIN2 previously cloned in pcDNA5 by Gibson) | F_AGCTGGGTCATAAACTGAATGACTACAAGGACCACGACGG<br><br>R_CCAGTGTGATGGATATCTGCCTATTAAGCTATCCCATCTCCTCC |
| pET-22-OSGIN1-$His_6$ | (Goupil et al., 2024) | - | - | - | - |
| pET-22-MBP-OSGIN1-$His_6$ | Gibson assembly | pET-22-MBP-HA-HsF30a-His<br><br>(Goupil et al., 2024) | F_ACCACCACCACCACCACTGATGAGATCCGGCTGCTAACAA<br><br>R_TGATCTTTCCGACTTGAACTAGCGTAATCTGGAACATCGT | pET-22b-Hs-OSGIN1<br><br>(Goupil et al., 2024) | F_ACGATGTTCCAGATTACGCTAGTTCAAGTCGGAAAGATCATCTCG<br><br>R_TTGTTAGCAGCCGGATCTCATCAGTGGTGGTGGTGGTGGT |
| pcDNA3.1(+) zeo-mNG2(1-10)-link-(11) | (Feng et al., 2017) (cloned in pcDNA | - | - | - | - |

| | | | | | |
|---|---|---|---|---|---|
| | also by Gibson- Gift from Michel Bouvier) | | | | |
| pcDNA5-mNG2(1-10)-3xFlag-OSGIN1 | Gibson assembly | pcDNA5-FTO-TO-TurboID-3xFlag-HsOSGIN1 (OSGIN1 previously cloned in pcDNA5 by Gibson) | F_CCGAGCTGAAGCACAGCATGCACGACATCGACTACAAGGA<br><br>R_TCTTCGCCCTTAGAAACCATTTAAGTTTAAACGCTAGAGTCC | pcDNA3.1(+)zeo-mNG2(1-10)-link-(11) | F_ACTCTAGCGTTTAAACTTAAATGGTTTCTAAGGGCGAAGAG<br><br>R_TCTTCGCCCTTAGAAACCATTTAAGTTTAAACGCTAGAGTCC |
| pcDNA5-mNG2(1-10)-3xFlag-OSGIN2 | Gibson assembly | pcDNA5-FTO-TO-UltraID-3xFlag-HsOSGIN2 (OSGIN2 was cloned in pcDNA5 by Gibson) | F_CCGAGCTGAAGCACAGCATGCACGACATCGACTACAAGGA<br><br>R_TCTTCGCCCTTAGAAACCATTTAAGTTTAAACGCTAGAGTCC | pcDNA3.1(+)zeo-mNG2(1-10)-link-(11) | F_ACTCTAGCGTTTAAACTTAAATGGTTTCTAAGGGCGAAGAG<br><br>R_TCCTTGTAGTCGATGTCGTGCATGCTGTGCTTCAGCTCGG |
| prK5-HA-OSGIN1-link-mNG2(11) | Gibson assembly | pRK5-HA-OSGIN1 (Goupil et al., 2024) | F_AAGCCTTCACCGACATGTGAGGCCCAACTTGTTTATTGCA<br><br>R_TACAGGGTACCGGCGGATCCCGGCGGTTTCCTAGTCTCTT | pcDNA3.1(+)zeo-mNG2(1-10)-link-(11) | F_AAGAGACTAGGAAACCGCCGGGATCCGCCGGTACCCTGTA<br><br>R_TGCAATAAACAAGTTGGGCCTCACATGTCGGTGAAGGCTT |
| prK5-HA-OSGIN2-link-mNG2(11) | Gibson assembly | pRK5-HA-OSGIN2 (OSGIN2 was previously cloned in pRK5 also by Gibson) | F_AAGCCTTCACCGACATGTGAGTCGACCTGCAGAAGCTTGG<br><br>R_TACAGGGTACCGGCGGATCCTTGAGCTCGAGATCTGAGTC | pcDNA3.1(+)zeo-mNG2(1-10)-link-(11) | F_GACTCAGATCTCGAGCTCAAGGATCCGCCGGTACCCTGTA<br><br>R_CCAAGCTTCTGCAGGTCGACTCACATGTCGGTGAAGGCTT |
| prK5-HA-link-mNG2(11) | Gibson assembly | pRK5-HA-OSGIN2 (OSGIN2 was previously cloned in pRK5 also by Gibson) | F_AAGCCTTCACCGACATGTGAGTCGACCTGCAGAAGCTTGG<br><br>R_TACAGGGTACCGGCGGATCCTTGAGCTCGAGATCTGAGTC | pcDNA3.1(+)zeo-mNG2(1-10)-link-(11) | F_GACTCAGATCTCGAGCTCAAGGATCCGCCGGTACCCTGTA<br><br>R_CCAAGCTTCTGCAGGTCGACTCACATGTCGGTGAAGGCTT |
| pCMV-Cit-Tuba1B-Kan/Neo | Restriction enzyme digestion and ligation | pCMV-Kan/Neo (pEGFP-N1 digested BamHI and Not I to remove EGFP and keep the empty backbone with Kanamycin and Neomicin/G418 selection cassettes) | XhoI and SacII restriction site digestion. | MSCV-CITRINE-Tuba1B-pgk-Puro (gift from Guy Sauvageau, Université de Montréal) | XhoI and SacII restriction site digestion of Citrine-Tuba1B |

### Table S2. Oligonucleotide sequences

| Purpose | Reference | Oligos (5'-3') |
|---|---|---|
| Sequencing OSGIN1 gDNA | (Goupil et al., 2024) | CTGATATTAGGGTAGGGATGTAGACCAGGC<br>GAGAATGAGGAAAGGGAGATGCCAGAGTC |
| sequencingOSGIN2 gDNA | | GGCCTTCAAGTTGGTATAGGTTGACATGTAG<br>GCTAAAATCAGATAAACCAATTACTCTCCAAG |
| qPCR for OSGIN1 | (Goupil et al., 2024) | TGCTGCAGAGGAAGCTCC<br>TCGGACAGGTAGTCCAGTC |
| qPCR for ACTB | Goupil et al., 2024) | ATTGGCAATGAGCGGTTC<br>TGAAGGTAGTTTCGTGGATGC |
| qPCR for GAPDH | Goupil et al., 2024) | AGCCACATCGCTCAGACAC<br>GCCCAATACGACCAAATCC |
